# Spatacsin regulates directionality of lysosome trafficking

**DOI:** 10.1101/2022.06.17.496577

**Authors:** Alexandre Pierga, Raphaël Matusiak, Margaux Cauhapé, Julien Branchu, Maxime Boutry, Frédéric Darios

**Author notes:** **Correspondence**: Frédéric Darios, Paris Brain Institute, INSERM U1127, CNRS UMR 7225, UPMC UMR S 1127, Hôpital de la Salpêtrière, 47, boulevard de l’Hôpital, F-75013 France, (F.D.).

## Abstract

The endoplasmic reticulum (ER) forms contacts with the lysosomal compartment, regulating lysosome positioning and motility. The movement of lysosomes are controlled by the attachment of molecular motors to their surface. However, the molecular mechanisms by which ER controls lysosome dynamics are still elusive. Here, we demonstrate that spatacsin is an ER-resident protein that regulates ER-lysosomes contacts to promote lysosome motility, shown by the presence of tubular lysosomes. Tubular lysosomes, which are highly dynamic, are entangled in a network of tubular ER. Screening for spatacsin partners required for tubular lysosome formation showed spatacsin to act by regulating protein degradation. We demonstrate that spatacsin promotes the degradation of its partner AP5Z1, which regulates the relative amount of spastizin and AP5Z1 at lysosomes. Spastizin and AP5Z1 contribute to lysosome trafficking by interacting with anterograde and retrograde motor proteins, kinesin KIF13A and dynein/dynactin subunit p150^Glued^, respectively. Ultimately, investigations in polarized neurons demonstrated that spatacsin-regulated degradation of AP5Z1 controls the directionality of lysosomes trafficking. Collectively, our results identify spatacsin as a protein regulating the directionality of lysosome trafficking.

## Introduction

Lysosomes are membrane-limited organelles responsible for the degradation of various cellular substrates. They degrade the content of late endosomes and autophagosomes upon fusion with these subcellular compartments. In addition, they also participate in many other cellular functions, such as cell metabolism and the repair of plasma membranes, as well as adhesion and migration (Ballabio and Bonifacino, 2020). These diverse functions rely on the cellular localization of lysosomes, as well as their motility and remodeling (Hipolito et al., 2018; Pu et al., 2016). Accordingly, lysosomes are a highly dynamic subcellular compartment (Bonifacino and Neefjes, 2017). They are retrogradely transported along microtubules upon coupling to cytoplasmic dynein and move anterogradely toward the cell periphery upon coupling to various kinesins (Ballabio and Bonifacino, 2020), changing their cellular distribution. The coordination of this bidirectional transport is particularly important for polarized cells such as neurons. However, the mechanisms regulating the coordination of anterograde or retrograde transports of lysosomes are not clearly established (Roney et al., 2022).

It has recently emerged that endosomes and lysosomes not only interact with the cytoskeleton but also form functional contacts with other subcellular organelles, in particular, the endoplasmic reticulum (ER). Such contacts with the ER are involved in the filling of lysosomes with Ca^2+^ or the non-vesicular transfer of lipids between the two subcellular compartments (Wang et al., 2017; Wilhelm et al., 2017). The interactions of the ER with endosomes and lysosomes also regulate the morphology and trafficking of these subcellular compartments. For example, the interaction of endosomes and lysosomes with the ER controls ER architecture by modulating the formation of the ER network at the cell periphery (Spits et al., 2021). Conversely, the ER regulates the distribution of endolysosomes through various mechanisms involving the proteins RNF26, protrudin, and ORP1L (Jongsma et al., 2016; Raiborg et al., 2015; Rocha et al., 2009) or it can regulate the morphology of endolysosomes by promoting their fission (Allison et al., 2017; Rowland et al., 2014). However, the control of endolysosomal dynamics is still only partially understood (Cabukusta and Neefjes, 2018) and the molecular mechanisms regulating lysosome dynamics at the levels of the ER have not been elucidated.

Lysosome function is impaired in various pathological conditions, such as, in neurodegenerative diseases (Oyarzún et al., 2019). Among them is hereditary spastic paraplegia type SPG11, which is due to loss-of-function mutations in the *SPG11* gene, leading to the absence of spatacsin (Stevanin et al., 2007). The subcellular localization of spatacsin is still debated, as it has been proposed to be localized at the ER, microtubules, or lysosomes (Hirst et al., 2013; Murmu et al., 2011). However, the loss of spatacsin function has been shown to impair lysosome function and distribution (Boutry et al., 2019; Branchu et al., 2017; Chang et al., 2014; Renvoisé et al., 2014; Varga et al., 2015), suggesting a lysosomal function for this protein. Spatacsin bears a Spatacsin_C domain in its C-terminus, which has been conserved throughout evolution up to plants (Patto and O’Kane, 2020). However, this domain has no homology in the human genome, suggesting a specific function. Spatacsin interacts with spastizin and AP5Z1, two proteins encoded by genes mutated in other forms of hereditary spastic paraplegia, SPG15 and SPG48, respectively (Hanein *et al*, 2008; Słabicki *et al*, 2010). Spastizin contains a FYVE domain, which binds to phosphatidylinositol-3-phosphate, allowing its recruitment to lysosomes (Hirst et al., 2021). AP5Z1 is a subunit of the adaptor protein complex AP5, involved in the sorting of proteins in late endosomes (Hirst et al., 2018). Loss-of-function mutations in *SPG11, SPG15*, or *SPG48* lead to the lysosomal accumulation of material (Branchu et al., 2017; Khundadze et al., 2019, 2013; Varga et al., 2015). However, it is not known how the absence of these proteins leads to lysosomal dysfunction and the mechanisms that regulate the interactions between these proteins have not been investigated.

Here, we show that spatacsin is an ER protein that regulates ER-lysosomes contacts to promote the motility of lysosomes in a ubiquitin-dependent manner. We show that spatacsin mediates the degradation of AP5Z1, which regulates the relative amounts of AP5Z1 at lysosomes. Moreover, by modulating AP5Z1 levels, spatacsin also indirectly regulates spastizin recruitment at lysosomes. As AP5Z1 and spastizin interact with motor proteins, the dynein/dynactin subunit p150^Glued^ and the kinesin KIF13A respectively, spatacsin is likely to regulate lysosome motility. Investigations in polarized axons demonstrate that spatacsin indeed regulates the directionality of lysosomes trafficking.

## Results

### Spatacsin is an endoplasmic reticulum-associated protein promoting ER-lysosomes contacts

We investigated the subcellular localization of spatacsin by transfecting mouse embryonic fibroblasts (MEFs) with a vector allowing the expression of spatacsin with a C-terminal V5 tag and determined its localization by immunostaining and confocal imaging. Spatacsin-V5 showed a diffuse distribution that poorly colocalized with late endosomes and lysosomes stained with Lamp1 (Figure 1A). By contrast, spatacsin-V5 partially colocalized with the ER labelled by the expression of GFP-Sec61β(Figure 1A). Consistently, spatacsin-V5 appeared to be mainly associated with the ER by STED imaging (Figure 1B). To confirm the subcellular localization of endogenous spatacsin, we used samples obtained from *Spg11* knockout mice (*Spg11*^-/-^) devoid of spatacsin and compared them to samples of wild-type mice (*Spg11*^+/+^). We prepared lysosomes and ER enriched fractions from *Spg11*^+/+^ and *Spg11*^-/-^ mouse brains using density gradients. Spatacsin was detected at very low levels in the lysosomal fraction, but the signal was stronger in the ER fraction, indicating that endogenous spatacsin is enriched in the ER (Figure 1C).

**Figure 1.**
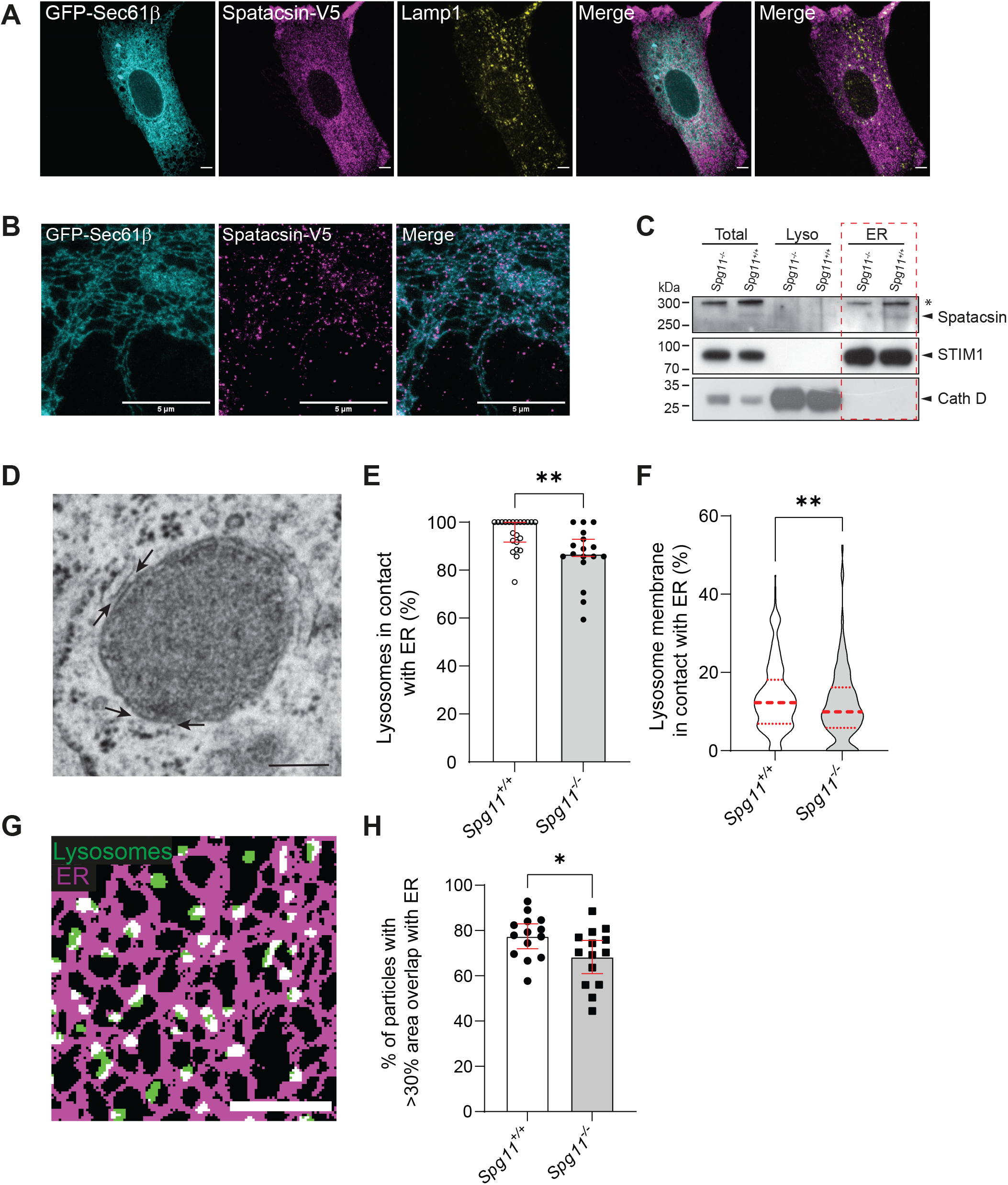
Spatacsin is an endoplasmic reticulum protein promoting ER-lysosomes contacts. A. MEFs expressing the ER marker GFP-Sec61β and V5-tagged spatacsin. Cells were immunostained with anti-V5 antibody and the lysosome marker Lamp1. Scale bar: 5 μm. B. STED images of MEFs expressing the ER marker GFP-Sec61β and expressing V5-tagged spatacsin. Cells were immunostained with anti-V5 and anti-GFP antibodies. Scale bar: 5 μm. C. Western blot analysis of ER and lysosome-enriched fractions obtained from *Spg11*^+/+^and *Spg11*^-/-^ mouse brains. Immunoblots with antibodies raised against spatacsin, ER protein STIM1, and lysosomal protein cathepsin D. ER fractions are encircled by the red rectangle. The asterisk indicates nonspecific signal. D. Transmission electron microscopy image of ER-lysosomes contacts in MEFs. Arrows point toward areas where ER and lysosomes are in contact. Scale bar: 200 nm. E. Proportion of lysosomes that make contact with the ER in transmission electron microscopy images in *Spg11*^+/+^ and *Spg11*^-/-^ MEFs. Median and 95% confidence interval (CI), N>17, cells from three independent experiments. **P<0.01, Mann-Whitney test. F. Quantification of the proportion of lysosomal membrane that is in contact with the ER in transmission electron microscopy images in *Spg11^+/+^* and *Spg11*^-/-^ MEFs. Median and quartiles, N=285 lysosomes from three independent experiments. **P<0.01, Mann-Whitney test. G. Binarized image of fibroblasts expressing GFP-Sec61β and Lamp1-mCherry. Note the overlap (white) between the ER (magenta) and lysosome (green) masks. Scale bar: 5 μm. H. Quantification of the proportion of lysosomes that have an area overlapping with the ER > 30% in *Spg11*^+/+^ and *Spg11*^-/-^ MEFs. Mean and 95% CI, N = 14 cells from three independent experiments. *P < 0.05, unpaired t-test.

The absence of spatacsin had no visible impact on the morphology of the ER, as observed by live confocal imaging (Supplementary Figure 1). As the loss of spatacsin function has been shown to affect lysosome function (Boutry et al., 2019, 2018; Branchu et al., 2017; Chang et al., 2014; Varga et al., 2015), we investigated if contacts between the ER and lysosomes were altered in absence of spatacsin (Figure 1D). Analysis of MEFs by transmission electron microscopy showed that more lysosomes were in contact with the ER in *Spg11*^+/+^ than in *Spg11*^-/-^ cells (Figure 1E). Moreover, the proportion of lysosomal membrane that was in contact with the ER was decreased in absence of spatacsin (Figure 1F). The decrease in contacts between the ER and lysosomes observed in absence of spatacsin was also detected by confocal microscopy. We transfected *Spg11*^+/+^ and *Spg11*^-/-^ MEFs with vectors expressing GFP-Sec61β, to label the ER, and Lamp1-mCherry, a marker of late endosomes and lysosomes (henceforth referred to as lysosomes). There was a decrease in the proportion of lysosomes with their area overlapping by more than 30% with the ER staining (Figures 1G and 1H). Overall, these data show that spatacsin is an ER-resident protein that promotes contacts between the ER and the lysosomes.

### Spatacsin regulates the dynamics of lysosomes that are moving along the ER

Since spatacsin promotes the formation of contacts between the ER and lysosomes that are important to modulate lysosome function (Bonifacino and Neefjes, 2017; Cabukusta and Neefjes, 2018), we evaluated how the loss of spatacsin may affect the lysosomes and their interaction with the ER.

Using MEFs transfected with a vector expressing Lamp1-mCherry, we observed by live imaging a higher number of lysosomes with tubular shape in *Spg11*^+/+^ compared to *Spg11*^-/-^ cells (Figures 2A-B), suggesting that spatacsin is required for the formation of tubular lysosomes. We also observed a higher number of tubular lysosomes in *Spg11*^+/+^ than *Spg11*^-/-^ MEFs when they were labelled with the acidic marker Lysotracker, with a pulse of Dextran-Texas Red followed by a long chase, or with DQ-BSA, which fluoresces upon degradation by lysosomal hydrolases (Marwaha and Sharma, 2017) (Supplementary Figures 2A-B). These results suggests that tubular lysosomes that are lost in absence of spatacsin represent a population of catalytically active lysosomes.

**Figure 2.**
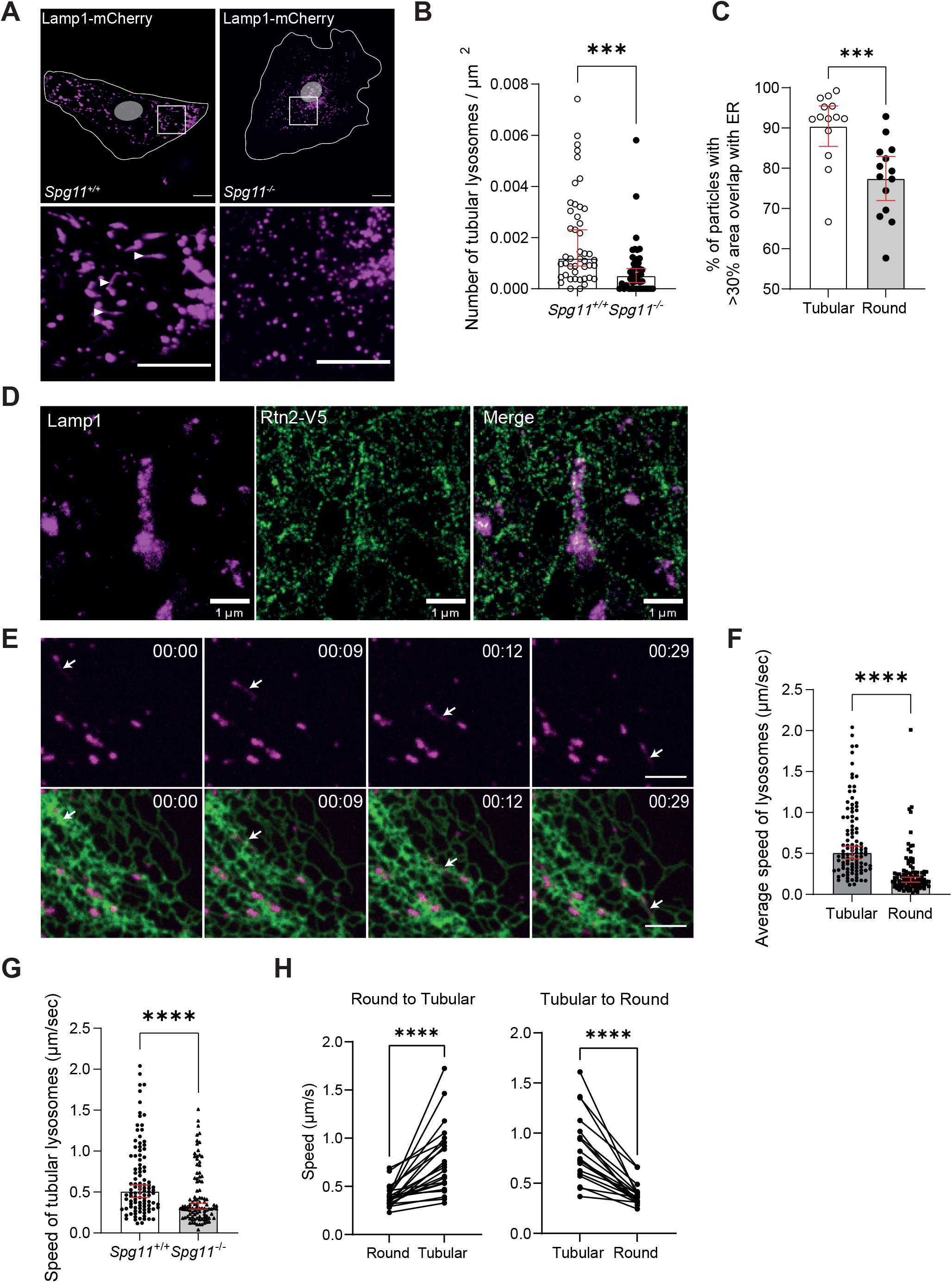
Spatacsin regulates the dynamics of lysosomes that are moving along the ER. A. Lamp1-mCherry expression in *Spg11*^+/+^ and *Spg11*^-/-^ MEFs imaged by spinning-disk confocal microscopy. Note the presence of tubular lysosomes in *Spg11*^+/+^ MEFs (white arrowheads). Scale bar: 5 μm. B. Quantification of the number of tubular lysosomes in *Spg11*^+/+^ and *Spg11*^-/-^ MEFs. Median and 95% CI, N = 45 cells from three independent experiments. ***P < 0.001. Mann-Whitney test. C. Quantification of the proportion of lysosomes that have an area overlapping with the ER > 30% based on their shape. Median and 95% CI, N = 14 cells from three independent experiments. ***P < 0.001, Mann Whitney test. D. STED image of a tubular lysosome stained with Lamp1 antibody and its close interaction with the ER tubular network stained by anti-V5 antibody targeting Reticulon2-V5 (Rtn2-V5) expressed in wild-type MEFs. Scale bar: 1 μm. E. Snapshot images of live imaging of a wild-type MEF transfected with Lamp1-mCherry (Magenta) and GFP-Sec-61β (Green). A tubular lysosome trafficking along the ER tubule network is indicated by an arrow. Scale bar: 5 μm. F. Average speed of tubular and round lysosomes in wild-type MEFs. Median and 95% CI, N = 110 from five independent MEFs. ****P < 0.0001, Mann-Whitney test. G. Quantification of the average speed of tubular lysosomes in *Spg11*^+/+^ and *Spg11*^-/-^ MEFs. Median and 95% CI, N > 100 lysosomes from five independent MEFs. ****P < 0.0001, Mann-Whitney test. H. Quantification of the instant speed of lysosomes in *Spg11*^+/+^ during their transition from round to tubular or tubular to round shape. N = 23 pairs of lysosomes from twelve independent MEFs. ****P<0.0001, Wilcoxon matched-pairs test.

To further investigate the properties of tubular lysosomes, we analyzed live images of MEFs expressing Lamp1-mCherry and GFP-Sec61β. This showed that the overlap of lysosome and ER staining was greater for tubular lysosomes than round lysosomes, suggesting that tubular lysosomes were in closer proximity to the ER (Figure 2C). Accordingly, STED microscopy showed that the tubular lysosomes were entangled in a network of ER tubules (Figure 2D). Live imaging also showed that tubular lysosomes moved along the ER tubular network (Figure 2E, Supplementary Video 1), suggesting that tubular lysosomes are closely associated with the ER network.

Since spatacsin promotes contacts between ER and lysosomes and is required for the formation of tubular lysosomes, we evaluated the impact of spatacsin on the properties of tubular lysosomes. Tracking individual lysosomes in wild-type MEFs by live imaging showed tubular lysosomes to move, on average, faster than round lysosomes (Figure 2F). The proportion of lysosomes with a speed > 0.3 μm/sec, corresponding to microtubule-dependent movement for these organelles (Cordonnier et al., 2001), was higher among the tubular than round lysosomes (Supplementary Figure 2C), suggesting that tubular lysosomes are highly mobile and dynamic. Comparison of the speed of lysosomes in *Spg11*^+/+^ and *Spg11*^-/-^ MEFs showed that tubular lysosomes move faster in *Spg11*^+/+^ than *Spg11*^-/-^ MEFs (Figure 2G, Supplementary Videos 2-3), whereas the dynamics of round lysosomes was not affected (Supplementary Figure 2D). Finally, tracking of individual lysosomes showed that lysosomes can change morphology from round to tubular. The transition between the two states was associated with a strong change in the speed of displacement (Figure 2H). The proportion of tubular lysosomes is thus an indicator of lysosome dynamics and was used as a metric to evaluate lysosome dynamics in the following experiments.

Overall, these data suggest that spatacsin regulates the formation and trafficking of tubular lysosomes that represent a population of catalytically active and highly dynamic lysosomes moving in close contact with the ER. As spatacsin is important for the formation and motility of tubular lysosomes, we next aimed to investigate how it could regulate their dynamics.

### Spatacsin regulates the dynamics of lysosomes by interacting with proteins involved in degradation pathways

We used MEFs derived from a mouse line in which exons 32 to 34 of *Spg11* are spliced out (Supplementary Figures 3A-C) to elucidate the molecular mechanisms by which spatacsin controls these lysosomal phenotypes. Such splicing retained the reading frame and led to the expression of a protein called spatacsin^Δ32-34^, which lacks a domain of 170 amino acids, partially deleting the conserved Spatacsin_C domain (Figure 3A). Western blot analysis of brains obtained from *Spg11*^+/+^, Spg*11*^-/-^, and *Spg11^Δ32-34/Δ32-34^* mice showed the latter strain to express a spatacsin protein that is slightly smaller than the wild-type protein (Supplementary Figure 3D).

**Figure 3.**
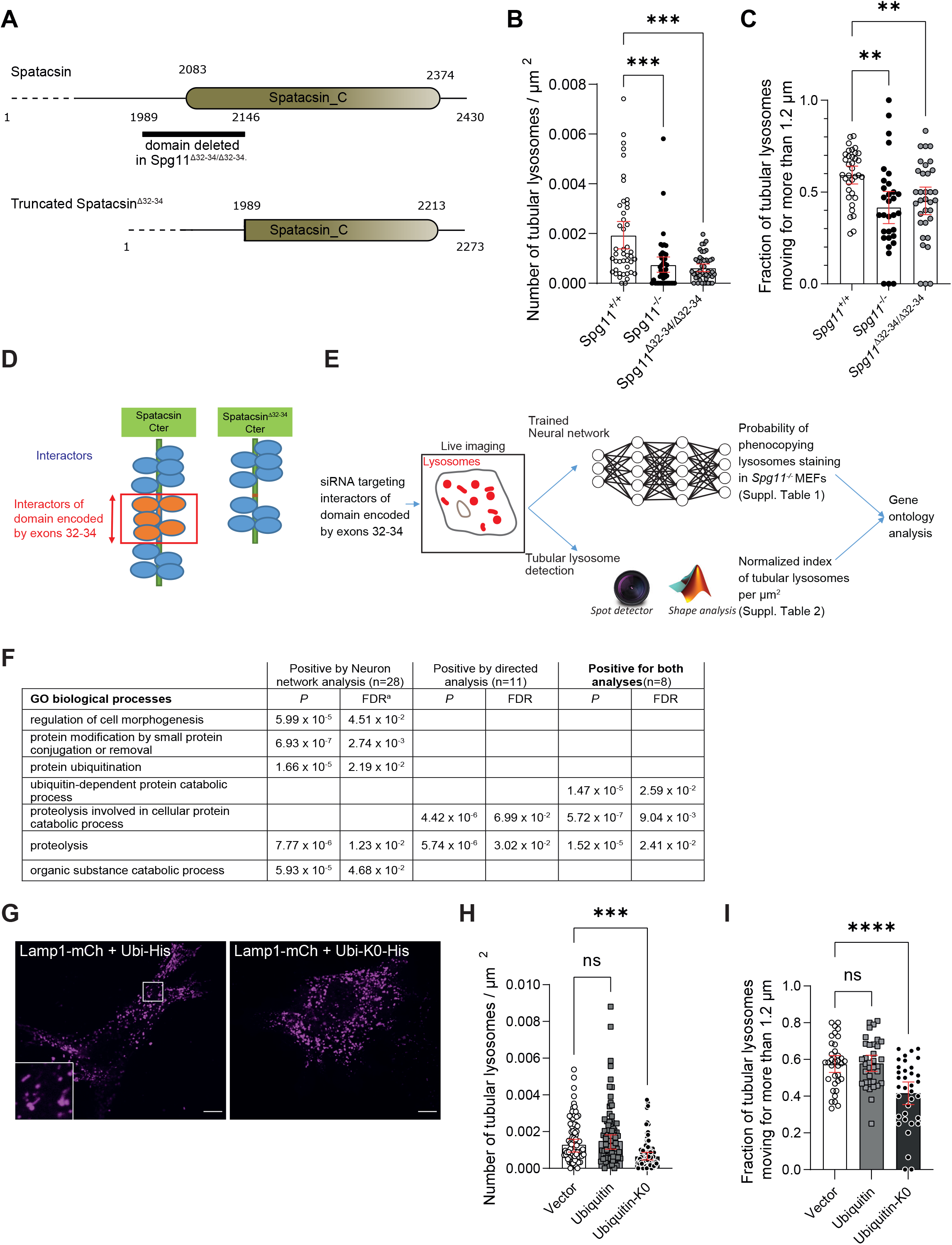
Spatacsin regulates lysosome dynamics by interacting with proteins involved in ubiquitin-dependent degradation. A. Representation of the Spatacsin_C domain and the region truncated in the *Spg11^Δ32-34/Δ32-34^* mouse model. B. Quantification of the number tubular lysosomes in MEFs devoid of spatacsin or expressing truncated spatacsin. Median and 95% CI, N=45 cells from 3 independent experiments. *** P<0.001 for both *Spg11*^-/-^ and *Spg1*^Δ*32-34*/Δ*32-34*^ when compared to *Spg11*^+/+^. Kruskall-Wallis test followed by Dunn’s multiple comparisons test. C. Quantification of the proportion of tubular lysosomes moving > 1.2 μm over 1 min in *Spg11*^+/+^, *Spg11*^-/-^, and *Spg11*^Δ*32-34*/Δ*32-34*^ MEFs. Mean and 95% CI, N = 35 cells from three independent experiments. **P < 0.01 for both *Spg11*^-/-^ and *Spg11*^Δ*32-34*/Δ*32-34*^ when compared to *Spg11*^+/+^ MEFs. One-way ANOVA followed by Sidak’s multiple comparisons test. D. Scheme showing the strategy to identify interactors of the domain of spatacsin encoded by exons 32 to 34 of *SPG11*. Yeast two-hybrid screens were performed with the C-terminal domain of human spatacsin (aa 1943-2443) or the same domain missing exons 32 to 34 as bait. Interactors specifically interacting with the spatacsin domain encoded by exons 32 to 34 (orange) were selected. E. Design of the screening process for interactors of the spatacsin domain encoded by exons 32 to 34 of *SPG11*. Each interactor was downregulated by siRNA in wild-type MEFs, and lysosomes were imaged by spinning disk confocal microscopy. The effect of the siRNAs was analyzed using an unbiased method (trained neural network) or directed analysis to quantify the presence of tubular lysosomes. F. Table summarizing the pathways identified by gene ontology analysis as being significantly enriched in the list of genes identified by neural network analysis, directed analysis, or both. FDR: False discovery rate. G. Live images of lysosomes in wild-type MEFs co-transfected with Lamp1-mCherry and either vector expressing ubiquitin (Ubi-His), or lysine-less ubiquitin-K0 (Ubi-K0-His). Scale bar: 10 μm. H. Quantification of the number of tubular lysosomes in wild-type MEFs transfected with an empty vector or vectors expressing ubiquitin or mutant ubiquitin-K0. Median and 95% CI, N = 75 cells from three independent experiments. ***P < 0.001, Kruskall-Wallis test followed by Dunn’s multiple comparisons test. I. Quantification of the proportion of tubular lysosomes moving > 1.2 μm over 1 min in wild-type MEFs transfected with an empty vector or vectors expressing ubiquitin or mutant ubiquitin-K0. Mean and 95% CI, N = 35 cells from three independent experiments. ****P < 0.0001, One-way ANOVA followed by Sidak’s multiple comparisons test.

Overexpression of spatacsin ^Δ32-34^ with a C-terminal V5 tag in MEFs showed diffuse and ER-associated localization like that of full-length spatacsin (Supplementary Figure 3E). Like *Spg11*^-/-^ MEFs, *Spg11^Δ32-34/Δ32-34^* fibroblasts had fewer tubular lysosomes than wild-type cells and had altered tubular lysosomes dynamics (Figures 3B-C). The difference in the dynamics of tubular lysosomes was validated by automated tracking, which showed tubular lysosomes to travel longer distance during one minute in *Spg11*^+/+^ than *Spg11*^-/-^ and *Spg11^Δ32-34/Δ32-34^* MEFs (Figure 3C). We used this method to analyze lysosomal dynamics in subsequent experiments. Overall, these results suggest that the spatacsin domain encoded by exons 32 to 34 of *Spg11* plays an important role in the formation and dynamics of tubular lysosomes.

We next aimed to define the molecular action of spatacsin in the formation of tubular lysosomes. We thus sought to identify proteins that bind to the domain encoded by exons 32 to 34 of *Spg11*. We performed a two-hybrid screen with the C-terminal region of human spatacsin (aa 1943-2443, containing the Spatacsin_C domain) and a second screen with the same C-terminal fragment in which the amino acids encoded by exons 32-34 were deleted. Comparison of the two screens identified several proteins that potentially bind directly to the domain encoded by exons 32 to 34 (Figure 3D, Supplementary Tables 1 and 2).

Among the proteins that could potentially bind to the domain encoded by exons 32 to 34, we aimed to identify those important for the regulation of lysosome dynamics. Thus, we downregulated each identified partner in wild-type MEFs using siRNA and analyzed the consequences on lysosomes, which were imaged by spinning disk confocal microscopy after staining with Dextran-Texas Red.

We used two methods to quantify the effect of siRNA on lysosomes (Figure 3E). First, we developed an unbiased classification method to discriminate between lysosomal staining in *Spg11*^-/-^ and *Spg11*^+/+^ MEFs by training a neural network that exploited all parameters of the lysosomal staining in images. The trained neural network was then used to predict the probability of the cell to be considered as a *Spg11*^-/-^ fibroblast for each image of fibroblasts transfected with siRNA. In parallel, we performed a directed analysis that automatically detected tubular lysosomes. For both methods, we evaluated how well the downregulation of each candidate using siRNA in wild-type MEFs phenocopied the lysosomal phenotype of *Spg11*^-/-^ MEFs. We compared the effect of each siRNA with that of three independent siRNAs downregulating *Spg11* (Supplementary Figure 3F). The neural network approach identified 28 genes and directed analysis identified 11 genes that, upon downregulation by siRNA, were at least as effective as *Spg11* siRNA to phenocopy *Spg11*^-/-^ MEFs (Figure 3E, Supplementary Tables 1 and 2). Eight genes were identified by both analyses, suggesting their importance in the function of spatacsin in lysosomes.

Gene ontology analysis of the candidates identified by the two approaches suggested a role of the ubiquitin-dependent protein catabolic process and proteolysis in modulation of the lysosomal phenotype (Figure 3F), suggesting that the action of spatacsin on lysosomes is linked to ubiquitin-dependent proteolysis. We confirmed this hypothesis by expressing mutant ubiquitin-K0, which prevents poly-ubiquitination of substrates required for degradation (Wu-Baer et al., 2003). Expression of this mutant in wild-type MEFs decreased the number of tubular lysosomes, as well as their dynamics (Figures3 G-I), suggesting that poly-ubiquitination of substrates and the promotion of ubiquitinated substrate degradation is necessary for the promotion of lysosomal dynamics by spatacsin.

### Spatacsin promotes UBR4-dependent degradation of AP5Z1

Our screening approach showed that the control of tubular lysosome formation and dynamics by spatacsin relies on a degradation pathway. However, exploration of the interactome of the spatacsin domain encoded by exons 32 to 34 revealed no binding partners with endolysosomal localization, suggesting that the degradation-dependent regulation may act on other proteins present in the endolysosomal system. Importantly, two partners of spatacsin, spastizin and AP5Z1, colocalize with lysosomes (Hirst et al., 2013).

We first investigated whether spastizin or AP5Z1 might be degraded in a spatacsin-dependent manner. We observed that overexpression of spatacsin-GFP in wild-type MEFs lowered levels of AP5Z1 while levels of spastizin where unaffected (Figure 4A). When polyubiquitination was blocked, AP5Z1 degradation by spatacsin was prevented (Figure 4B), indicating that this degradation likely relies on spatacsin ability to interact with its partners involved in protein degradation. Accordingly, the degradation of AP5Z1 was not observed upon overexpression of spatacsin^Δ32-34^-GFP, suggesting that the domain encoded by exons 32 to 34 contains the information that controls the degradation of AP5Z1 (Figure 4C). This domain of spatacsin notably interacts with UBR4. We confirmed by co-immunoprecipitation that UBR4 interacts with the C-terminal domain of spatacsin, but not the domain lacking the fragment encoded by exons 32 to 34 of *Spg11* (Figure 4D). Downregulation of UBR4 prevented the degradation of AP5Z1 mediated by spatacsin-GFP overexpression, suggesting that spatacsin mediates UBR4-dependent degradation of AP5Z1 (Figure 4A).

**Figure 4.**
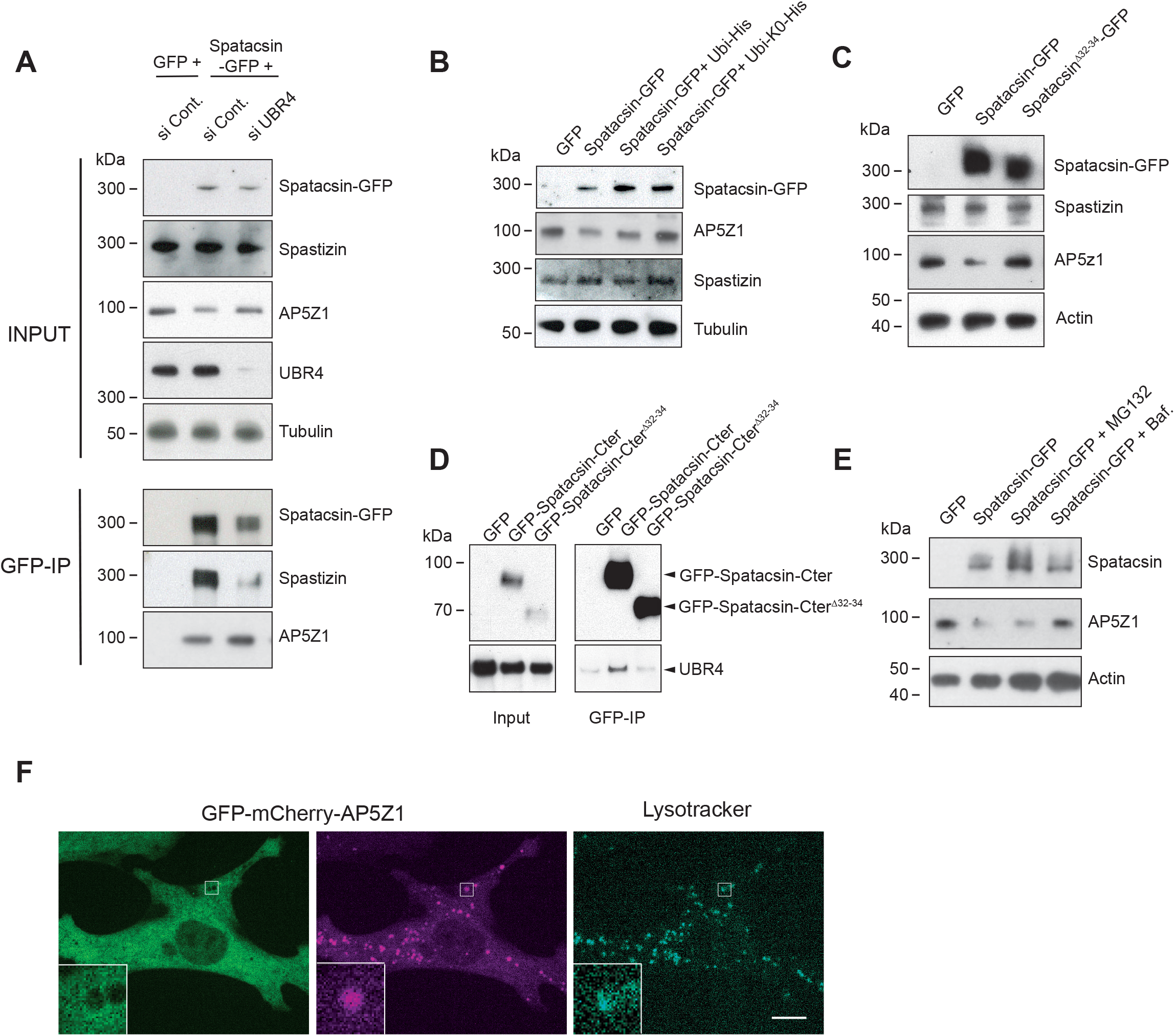
Spatacsin promotes UBR4-dependent degradation of AP5Z1. A. Western blot showing lysates and GFP immunoprecipitations in wild-type MEFs transfected with vectors expressing GFP, or spatacsin-GFP with control siRNA or siRNA downregulating UBR4. Top panel: inputs showing that overexpression of spatacsin-GFP leads to lower levels of AP5Z1, an effect that is prevented upon UBR4 downregulation. Bottom: GFP-immunoprecipitation showing that downregulation of UBR4 increases the co-immunoprecipitation of AP5Z1 with spatacsin-GFP and decreases the co-immunoprecipitation of spastizin with spatacsin-GFP. B. Western blots showing that downregulation of AP5Z1 levels upon overexpression of spatacsin-GFP is inhibited by co-expression of mutant ubiquitin-K0. C. Western blots showing levels of spatacsin-GFP, spastizin, AP5Z1, and actin in MEFs overexpressing GFP, spatacsin-GFP, or spatacsin^Δ32-34^-GFP. Note that overexpression of full-length spatacsin-GFP, but not spatacsin^Δ32-34^-GFP, leads to lower levels of endogenous AP5Z1. D. Western blots showing co-immunoprecipitation of UBR4 with the C-terminal domain of spatacsin (aa 1943-2443, GFP-spatacsin-Cter) but not the construct lacking amino acids encoded by exons 32 to 34 (GFP-spatacsin-Cter^Δ32-34^). E. Western blot monitoring expression levels of AP5Z1 in wild-type MEFs overexpressing spatacsin-GFP and treated for 16 hours with either MG132 (15μM) or bafilomycin (Baf., 100 nM). Actin immunoblot was used as loading control. Note that bafilomycin treatment leads to normal levels of AP5Z1 upon overexpression of spatacsin-GFP. F. Live images of wild-type MEFs expressing GFP-mCherry-AP5Z1 and stained with Lysotracker Blue. Inset shows mCherry-AP5Z1 colocalized with acidic lysosomes labelled by Lysotracker while GFP-AP5Z1 is located on the surface of lysosomes. Scale bar: 10 μm.

Furthermore, the decrease in AP5Z1 levels upon spatacsin-GFP overexpression was blocked when MEFs were treated with bafilomycin, but not with the proteasome inhibitor MG132 (Figure 4E), suggesting that AP5Z1 is degraded by lysosomes in a spatacsin-dependent manner. To test this hypothesis, we expressed AP5Z1 fused to both GFP and mCherry and analyzed the colocalization of fluorescence with the acidic lysosome marker, lysotracker (Figure 4F). In wild-type MEFs, mCherry was colocalized with lysosomes. In contrast, GFP that is sensitive to pH was poorly colocalized with lysosomes, suggesting that AP5Z1 was mainly inside the acidic subcellular compartment (Figure 4F). However, GFP was slightly enriched around lysosomes, suggesting that some AP5Z1 may be localized at the lysosomal surface. Of note, GFP signal was recovered when MEFs were treated with bafilomycin to neutralize lysosomal pH, confirming that most AP5Z1 was inside the acidic compartment (Supplementary Figure 4).

Together these results suggest that spatacsin contributes to the translocation of AP5Z1 inside lysosomes to promote its degradation in a UBR4-dependent manner.

### Spatacsin-mediated degradation of AP5Z1 promotes spastizin recruitment to lysosomes

Spatacsin also interacts with spastizin, and is required to recruit spastizin to lysosomes (Hirst et al., 2021). Accordingly, we observed weaker colocalization of spastizin-GFP with Lamp1-mCherry in *Spg11*^-/-^ than *Spg11*^+/+^ MEFs by live imaging (Figures 5A-B). We hypothesized that spatacsin interaction with spastizin was required for spastizin localization to lysosomes. To test this hypothesis, we performed a proximity-ligation assay in MEFs transfected with spatacsin-V5 and spastizin-HA. This assay showed that the interaction of spatacsin with spastizin occurred at lysosomes labelled by Lamp1 immunostaining (Figure 5C). This suggests that the spatacsin-spastizin interaction occurs at contact sites between the ER and lysosomes to allow spastizin recruitment to lysosomes.

**Figure 5.**
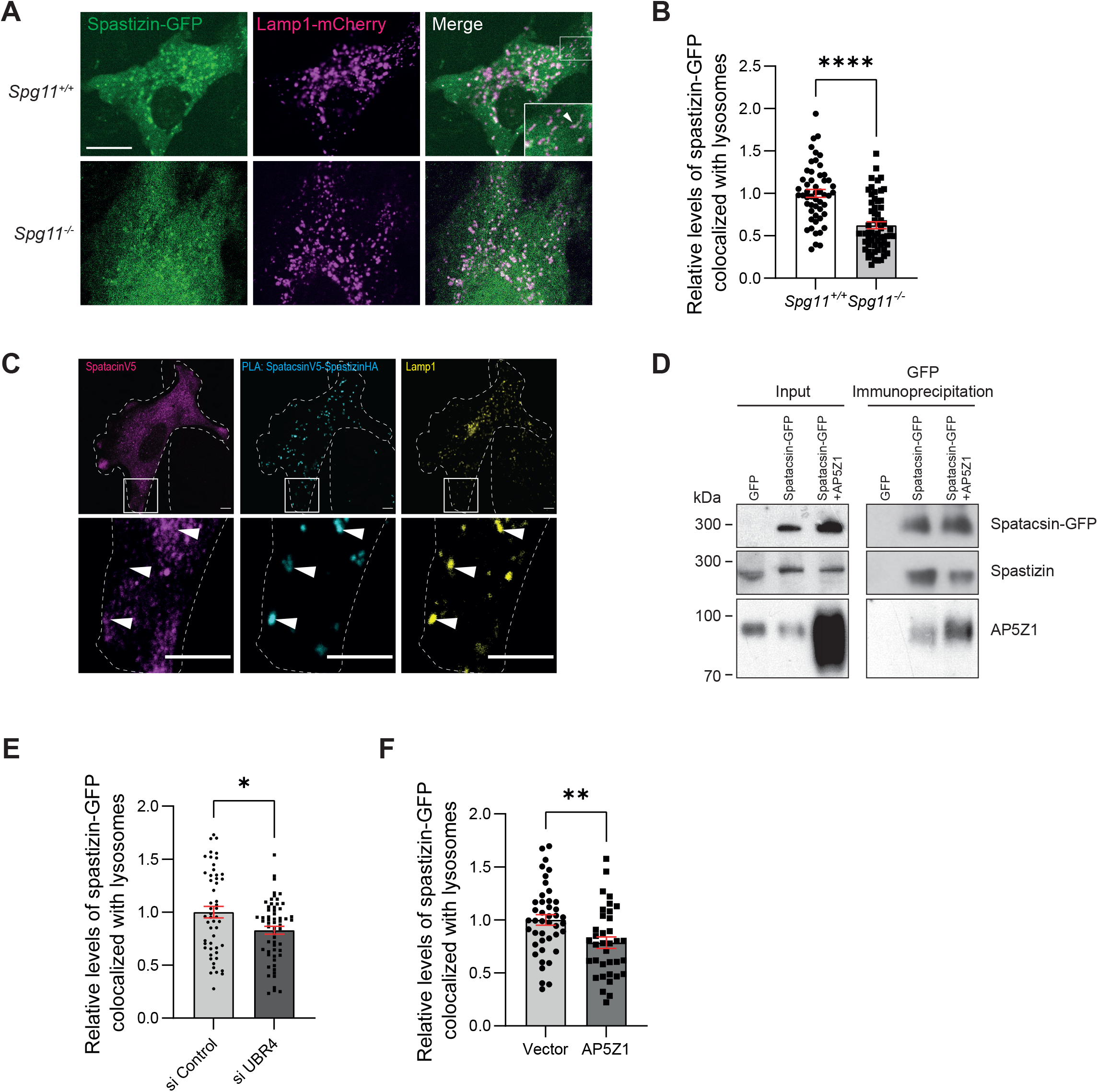
Spatacsin-mediated degradation of AP5Z1 promotes spastizin recruitment to lysosomes. A. Expression of spastizin-GFP and Lamp1-mCherry in *Spg11*^+/+^ and *Spg11*^-/-^ MEFs. Note the localization of spastizin-GFP along the tubular lysosomes in *Spg11*^+/+^ MEFs (arrowheads in insert), and the weaker colocalization of spastizin-GFP with lysosomes in *Spg11*^-/-^ MEFs. Scale bar 10 μm. B. Quantification of the proportion of spastizin-GFP colocalized with Lamp1-mCherry in *Spg11*^+/+^ and *Spg11*^-/-^ MEFs. Mean and 95% CI, N > 50 cells from three independent experiments. ****P < 0.0001, unpaired t-test. C. Proximity-ligation assay (PLA) showing the interaction between V5-tagged spatacsin and HA-tagged spastizin in wild-type MEFs. The PLA signal (cyan) is detected at the level of lysosomes immunostained with Lamp1 (arrowheads). Scale bar: 5 μm. D. Co-immunoprecipitation of spastizin and AP5Z1 with spatacsin-GFP. Note that the interaction of spatacsin-GFP with spastizin decreases when AP5Z1 is overexpressed. E. Quantification of the proportion of spastizin-GFP colocalized with Lamp1-mCherry in wild-type MEFs expressing siUBR4. Mean and 95% CI, N > 59 cells from three independent experiments. * P=0.0102, unpaired t-test. F. Quantification of the proportion of spastizin-GFP colocalized with Lamp1-mCherry in wild-type MEFs overexpressing AP5Z1. Mean and 95% CI, N > 36 cells from three independent experiments. **P < 0.01, unpaired t-test.

We then investigated whether the degradation of AP5Z1 mediated by spatacsin and UBR4 had an impact on the interaction of spatacsin with spastizin by coimmunoprecipitation. Downregulation of UBR4 that prevented degradation of AP5Z1 mediated by spatacsin (Figure 4A) led to higher interaction of spatacsin with AP5Z1 and decreased the interaction of spatacsin with spastizin (Figure 4A). This suggested that interaction of spatacsin with spastizin relies on the levels of AP5Z1. To test this hypothesis, we overexpressed AP5Z1. This overexpression lowered the amount of spastizin coimmunoprecipitated with spatacsin (Figure 5D), suggesting that AP5Z1 competes with spastizin to bind spatacsin. Both downregulation of UBR4 and overexpression of AP5Z1 partially prevented association of spastizin with lysosomes (Figures 5E-F), suggesting that decreasing spatacsin-spastizin interaction impaired spastizin recruitment to lysosomes.

Overall, these data suggest that AP5Z1 competes with spastizin to interact with spatacsin, and that AP5Z1 prevents association of spastizin with lysosomes.

### Spastizin and AP5Z1 regulate association of lysosomes with motor proteins

We then investigated whether AP5Z1 and spastizin regulate lysosome dynamics. To evaluate whether AP5Z1 levels impact lysosomes dynamics, we co-transfected wild-type MEFs with Lamp1-mCherry and a vector expressing GFP-AP5Z1. Overexpression of GFP-AP5Z1 decreased the number of tubular lysosomes (Figure 6A) and impaired lysosomal dynamics in wild-type MEFs (Figure 6B). To evaluate the role of spastizin at the lysosomes, we treated wild-type MEFs with the PI3 kinase inhibitor wortmannin that prevented the lysosomal localization of spastizin (Figure 6C). This treatment decreased the number of tubular lysosomes and their dynamics (Figure 6D-Supplementary Figure 5A). Therefore, both AP5Z1 and spastizin regulate lysosome motility.

**Figure 6.**
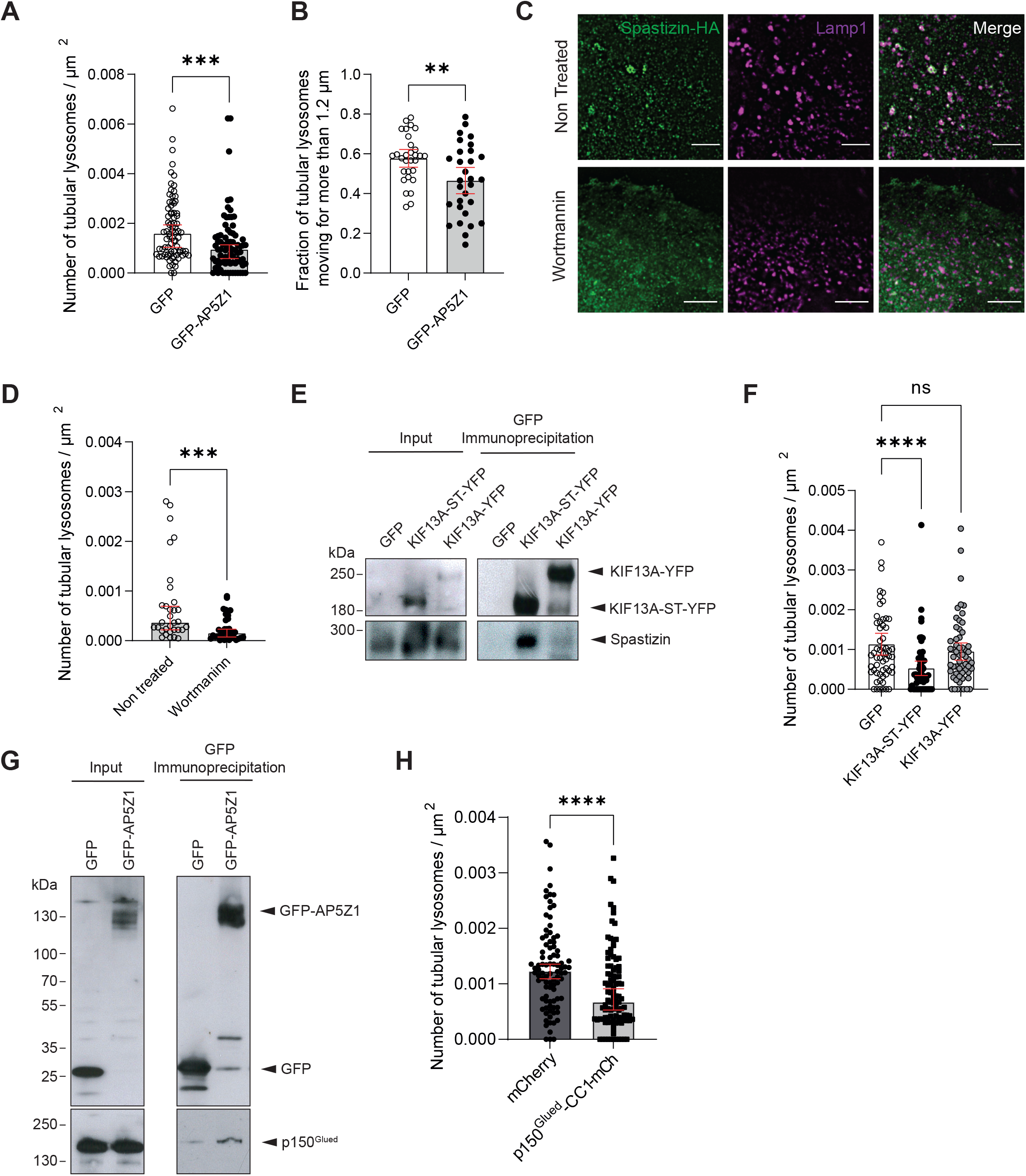
Spastizin and AP5Z1 regulate association of lysosomes with motor proteins. A. Quantification of the number of tubular lysosomes in wild-type MEFs transfected with a vector overexpressing GFP-AP5Z1. Median and 95% CI, N = 78 cells from three independent experiments. ***P < 0.001, Mann-Whitney test. B. Quantification of the proportion of tubular lysosomes moving > 1.2 μm over 1 min in wild-type MEFs transfected with a vector overexpressing GFP-AP5Z1. Mean and 95% CI, N = 30 cells from three independent experiments. **P < 0.01, unpaired t-test. C. Images of wild-type MEFs expressing spastizin-GFP and Lamp1-mCherry treated with 100 nM wortmannin for 1 h. Note the loss of colocalization of spastizin-GFP and Lamp1-mCherry upon wortmannin treatment. Scale bar: 5 μm. D. Quantification of the number of tubular lysosomes in wild-type MEFs treated with wortmannin. Median and 95% CI, N = 32 cells from three different independent experiments. ***P = 0.0009, Mann-Whitney test. E. Western blots showing co-immunoprecipitation of spastizin-HA with KIF13A-YFP or mutant KIF13A-ST-YFP (devoid of the motor domain). F. Quantification of the number of tubular lysosomes in wild-type MEFs transfected with wild-type KIF13A or mutant KIF13A-ST. Median and 95% CI, N = 58 cells from three different independent experiments. ****P < 0.0001, Kruskall-Wallis test. G. Western blots showing co-immunoprecipitation of p150^Glued^ with GFP-AP5Z1. H. Quantification of the number of tubular lysosomes in wild-type MEFs transfected with a vector overexpressing p150^Glued^-CC1-mCherry. Median and 95% CI, N = 90 cells from three independent experiments. ****P < 0.001, Mann-Whitney test.

Spastizin has been shown to interact with the motor protein KIF13A (Sagona et al., 2010). Overexpression of the mutant KIF13A-ST-YFP, devoid of the motor domain (Delevoye et al., 2014) but capable of interacting with spastizin (Figure 6E), prevented the formation of tubular lysosomes and altered their dynamics (Figure 6F-Supplementary Figure 5B). Overall, these results suggest that spastizin mediates the recruitment of KIF13A to lysosomes to control the formation of tubular lysosomes and their motility.

However, the formation of tubular membrane organelles requires the coordination of numerous effectors (Anitei and Hoflack, 2012). We investigated whether AP5Z1 might also interact with some motor protein. The endocytic adaptor protein complex AP2 was shown to interact with p150^Glued^, a subunit of dynein/dynactin complex implicated in retrograde transport (Kononenko et al., 2017; Waterman-Storer et al., 1997). Co-immunoprecipitation showed that AP5Z1 also interacted with p150^Glued^ (Figure 6G). To test whether this protein contributes to lysosome dynamics, we expressed a dominant negative construct p150^Glued^-CC1 (Quintyne et al., 1999), which prevented the formation of tubular lysosomes and altered their dynamics (Figure 6H-Supplementary Figure 5C). Overall, these results suggest that AP5Z1 mediates the recruitment of p150^Glued^ to lysosomes to control the formation of tubular lysosomes and their motility.

Together these data show that both spastizin and AP5Z1 contribute to lysosome trafficking by interacting with anterograde and retrograde motor proteins, respectively. Since spatacsin regulates the levels of AP5Z1 as well as the association of spastizin with lysosomes, the balance between spastizin and AP5Z1 may regulate the directionality of lysosome movements.

### Spatacsin regulates directionality of lysosome trafficking in axons

We then investigated whether spatacsin may regulate lysosome trafficking in highly polarized neurons that are degenerating in the absence of spatacsin, spastizin or AP5Z1 (Branchu et al., 2017; Khundadze et al., 2019, 2013; Varga et al., 2015).

Primary cultures of wild-type or *Spg11*^-/-^ neurons were transfected with Lamp1-GFP and mCherry-TRIM46 and analyzed after 7 days *in vitro*. Since the uniform polarity of microtubules in axons facilitates the analysis of transport (Yau et al., 2016), the analysis of lysosomal trafficking was focused on axons, which were identified by the presence of the axon initial segment protein TRIM46 (Supplementary Figure 6A). Like fibroblasts, neurons devoid of spatacsin presented a lower number of tubular lysosomes along the axon (Figures 7A-B). It also appeared that, along the axon, a lower proportion of particles were engaged in a directional movement in absence of spatacsin (Figures 7C-D). Furthermore, analysis of the directionality of movement showed a lower proportion of lysosomes with anterograde movement in *Spg11*^-/-^ axons compared to wild-type axons, whereas the proportion of lysosome with retrograde movement was not modified (Figure 7E-Supplementary Figure 6B). Importantly, axons of wild-type but not *Spg11*^-/-^ neurons presented similar proportions of lysosomes with anterograde and retrograde movement, (Figure 7E-Supplementary Figure 6B). The imbalance between anterograde and retrograde movement observed in *Spg11*^-/-^ axons led to a concentration of lysosomes in the proximal part of axons of *Spg11*^-/-^ neurons compared to wild-type neurons (Figure 7F). Together, these data suggest that spatacsin regulates the equilibrium between anterograde and retrograde trafficking of lysosomes and therefore affects the distribution of lysosomes along the axon.

**Figure 7.**
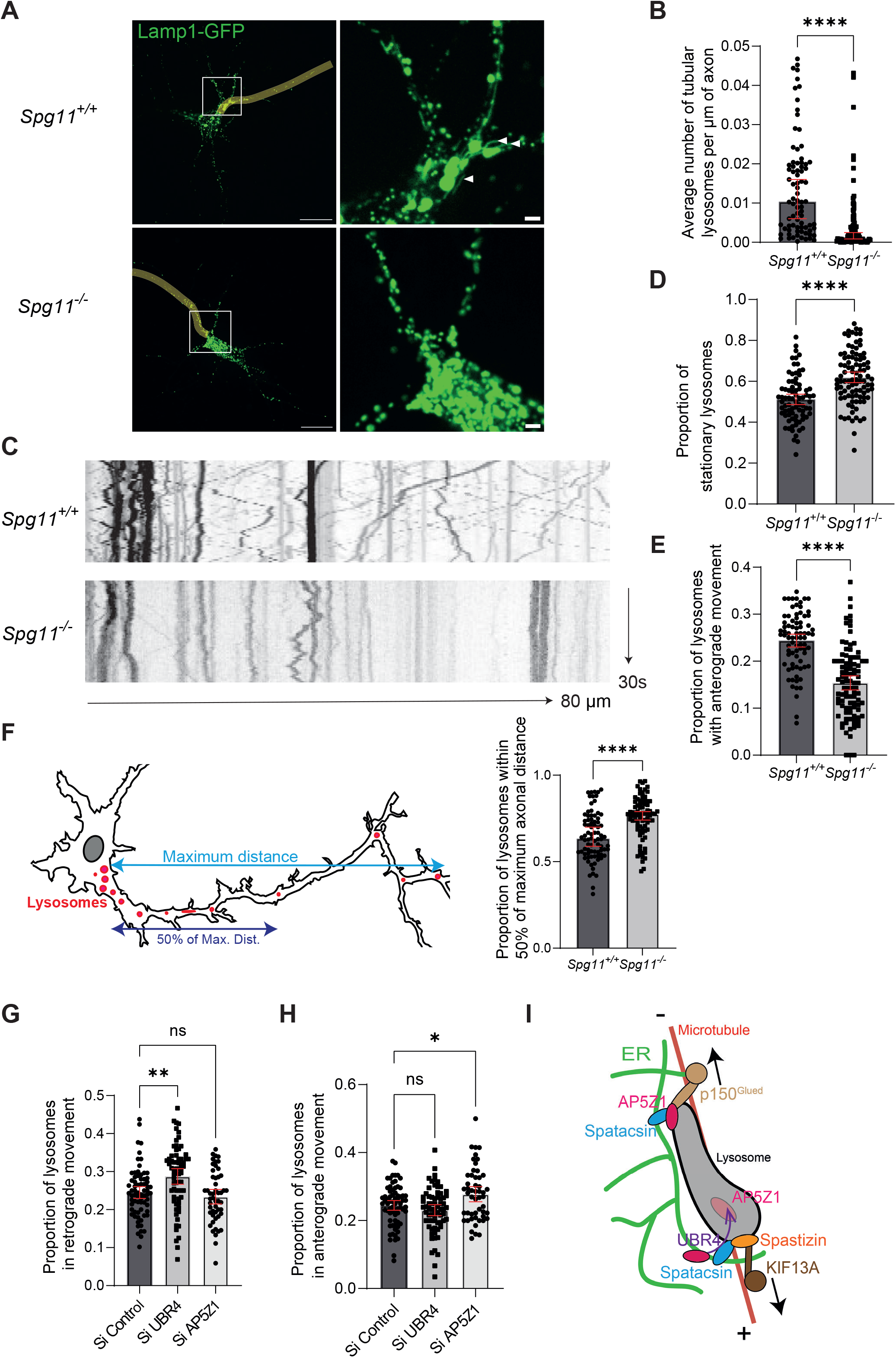
Spatacsin regulates directionality of lysosome trafficking in axons. A. Live images of *Spg11*^+/+^ and *Spg11*^-/-^ primary mouse neurons at DIV7 expressing Lamp1-GFP. Axons are highlighted in yellow. White arrows point tubular lysosomes. Note the loss of tubular lysosomes in *Spg11*^-/-^ neurons. Scale bar: 10 μm, inset: 1 μm. B. Quantification of the number of tubular lysosomes along the axon in *Spg11*^+/+^ and *Spg11*^-/-^ mouse neurons. Median and 95% CI, N = 80 cells from three independent experiments. ****P < 0.0001, Mann-Whitney test. C. Kymographs representing lysosomes movement over 30 seconds along the axon of *Spg11*^+/+^ and *Spg11*^-/-^ primary mouse neurons. Left side of the kymographs is toward the soma and right is toward the axon. Length of the axon: 80 μm. D. Quantification of the proportion of lysosomes that are stationary along the axon of *Spg11*^+/+^ and *Spg11*^-/-^ primary mouse neurons. Mean and 95% CI, N = 79 cells from three independent experiments. ****P < 0.0001, unpaired t-test. E. Quantification of the proportion of lysosomes that are moving anterogradely along the axon of *Spg11*^+/+^ and *Spg11*^-/-^ primary mouse neurons. Mean and 95% CI, N = 79 cells from three independent experiments. ****P < 0.0001, unpaired t-test. F. *Left:* Scheme representing the distribution of lysosomes along the axon of mouse neurons, the maximum distance considered is from the farthest detected axonal lysosome from the soma to the beginning of the axon. *Right:* Quantification of the proportion of lysosomes that are found within 50% of the maximum distance along the axon in *Spg11*^+/+^ and *Spg11*^-/-^ primary mouse neurons. Median and 95% CI, N = 80 cells from three independent experiments. ****P < 0.0001, Mann-Whitney test. G. Quantification of the proportion of lysosomes that are moving retrogradely along the axon of *Spg11*^+/+^ primary mouse neurons upon transfection with Lamp1-GFP and siRNA downregulating either UBR4 or AP5Z1. Mean and 95% CI, N = 70 cells for siControl & siUBR4, N=50 for siAP5Z1 from three independent experiments. *P = 0.017, one-way ANOVA followed by Sidak’s multiple comparisons test. H. Quantification of the proportion of lysosomes that are moving anterogradely along the axon of *Spg11^+/+^* primary mouse neurons upon transfection with Lamp1-GFP and siRNA downregulating either UBR4 or AP5Z1. Mean and 95% CI, N = 70 cells for siControl & siUBR4, N=50 for siAP5Z1from three independent experiments. **P < 0.01, one-way ANOVA followed by Sidak’s multiple comparisons test. I. Scheme showing that spatacsin regulates the levels of AP5Z1 by promoting its degradation in a UBR4-depedent manner. This regulates the amount of AP5Z1 and spastizin at the lysosome surface. Since AP5Z1 and spastizin interact with the retrograde p150^Glued^ and anterograde KIF13A motor proteins, respectively, the regulation of AP5Z1 and spastizin levels at lysosome surface is a mechanism regulating directionality of lysosome trafficking.

We then investigated whether the action of spatacsin on lysosome trafficking in axons may be regulated by the UBR4 mediated degradation of AP5Z1. Downregulation of UBR4 in neurons expressing Lamp1-GFP increased the proportion of lysosomes with retrograde movement (Figure 7G). Interestingly, over-expression of GFP-AP5Z1 in neurons expressing Lamp1-mCherry had similar consequences on promoting retrograde movement in axons (Supplementary Figures 6 C-D). Conversely, downregulation of AP5Z1 levels using an siRNA (Supplementary Figure 6E), promoted anterograde movement (Figure 7H). Overall, these results show that the levels of AP5Z1 are critical to control the directionality of lysosome trafficking in axons. As spatacsin regulates the levels of AP5Z1 by promoting its degradation by lysosomes in a UBR4-dependent manner, this mechanism likely explains the change in the directionality of lysosome trafficking and repartition of lysosomes in the axon of *Spg11*^-/-^ neurons (Figure 7I).

## Discussion

The loss of spatacsin, involved in hereditary spastic paraplegia type SPG11, causes lysosomal dysfunction (Boutry et al., 2018; Branchu et al., 2017; Chang et al., 2014; Varga et al., 2015). However, the molecular function of spatacsin has, thus far, remained elusive. Here, we establish that spatacsin is a protein localized at contacts between the ER and lysosomes to control the trafficking of lysosomes. Using a combination of trained neural network and targeted image analysis coupled to an siRNA screen, we show that spatacsin function is regulated by protein degradation pathways. We demonstrate that spatacsin controls the directionality of lysosome movement by modulating the degradation of AP5Z1.

Contacts between the ER and lysosomes are mediated by a variety of molecular actors, and play a role in numerous functions such as regulation of calcium homeostasis, cholesterol transfer between compartments or regulation of lysosomes positioning or dynamics (Wang et al., 2017; Wilhelm et al., 2017). Several molecular factors regulating contacts between the ER and lysosomes restrict lysosome dynamics. For example, SNX19 or the ubiquitination of p62 by the ER-localized ubiquitin ligase RNF26 maintain lysosomes in the perinuclear region of cells (Jongsma et al., 2016; Saric et al., 2021). When cholesterol content in lysosome membrane is low, the Rab7 effector ORP1L adopts a conformation allowing its interaction with the ER protein VAP, maintaining lysosomes in the cellular periphery (Rocha et al., 2009). In contrast, we show here that spatacsin is an ER-localized protein that promotes contacts between the ER and lysosomes and regulates lysosome motility, highlighting the diversity of functions controlled by contact sites between the ER and lysosomes.

The importance of spatacsin in lysosome function is revealed by its impact on tubular lysosomes. We demonstrate here that these lysosomes observed only by live imaging are present both in mouse embryonic fibroblasts and neurons. They are catalytically active lysosomes that are more mobile than punctate lysosomes as observed in other cell types (Mrakovic et al., 2012). The proportion of tubular lysosomes was thus used as an indicator of the general dynamic state of the lysosomal compartment. Consistently, all experimental conditions decreasing the proportion of tubular lysosomes also impaired their dynamics, suggesting that the formation of tubular lysosomes is associated with their trafficking. Formation of tubular lysosomes and lysosome trafficking were previously associated with the kinesin KIF5B (Du et al., 2016; Guardia et al., 2016). Here, we demonstrate the importance of another kinesin, KIF13A, or the dynein/dynactin subunit p150^Glued^. These motor proteins likely contribute to the formation of tubular lysosomes by pulling on membranes, as shown in endosomes (Delevoye et al., 2016; Hong et al., 2009).

The motor proteins KIF13A and p150^Glued^ are likely associated to lysosomes by spastizin and AP5Z1, respectively, which may be considered as motor adaptor molecules. Spastizin interacts with KIF13A, which is a plus end-directed microtubule motor that allows the movement of vesicles toward the cell periphery (Nakagawa et al., 2000). In contrast, AP5Z1 interaction with the subunit of dynein/dynactin p150^Glued^ likely promotes the movement of lysosomes toward the minus-end of microtubules (Waterman-Storer et al., 1997). Our data show that spatacsin regulates the association of spastizin and AP5Z1 with the membrane of lysosomes. The association of spastizin with lysosomes requires its interaction with spatacsin at the level of contact sites between the ER and lysosomes. Importantly, AP5Z1 appears to prevent the association of spastizin with lysosomes. The degradation of AP5Z1 mediated by spatacsin is thus a mechanism to control the relative amount of both spastizin and AP5Z1 at the lysosomal surface. Since AP5Z1 and spastizin interact with a minus-end or a plus-end-directed motor protein, respectively, spatacsin likely orientates the direction of lysosome movement by modulating the equilibrium between AP5Z1 and spastizin at the lysosome surface (Figure 7I).

This function of spatacsin is confirmed by the lower anterograde trafficking of lysosomes observed in axons of *Spg11*^-/-^ neurons. A similar phenotype was observed in neurons devoid of spastizin (Marrone et al., 2022). Furthermore, the coupling of motor proteins to lysosomes mediated by spatacsin may also contribute to the distribution of lysosomes in other cell types and may explain the clustering of lysosomes around the nucleus observed in fibroblasts when spatacsin or spastizin are absent (Boutry et al., 2019; Marrone et al., 2022).

The imbalance between retrograde and anterograde transport in *Spg11*^-/-^ axons resulted in the enrichment of lysosomes toward the neuronal soma while the axons were slightly deprived of lysosomes. Axonal lysosome availability has been shown to be regulated notably by ER-lysosome contact sites (Özkan et al., 2021). Spatacsin is present at these sites and its regulation of lysosomal repartition along the axon is concordant with this finding. Moreover, several recent papers have highlighted the importance of axonal lysosomal trafficking in the development of neurodegenerative disease (Lie and Nixon, 2019; Roney et al., 2022). In SPG11 models, previous work has shown that cultured neurons deprived of spatacsin presented axonal instability (Pérez-Brangulí et al., 2014). Therefore, the impaired distribution of lysosomes along the axon in absence of spatacsin which results of impaired lysosomal dynamics appears to be particularly relevant to the development of hereditary spastic paraplegia type 11.

The regulation of lysosome trafficking by spatacsin relies on the tight regulation of AP5Z1 levels. Spatacsin promotes its degradation by lysosomes. However, spatacsin required some partners, such as UBR4, to mediate AP5Z1 degradation. Interestingly, UBR4 associates with cellular cargoes destined to autophagic vacuoles and degraded by the lysosome (Kim et al., 2013), which is consistent with the inhibition of AP5Z1 degradation observed upon blockade of lysosome degradative activity by bafilomycin. Our screening strategy revealed several other proteins important for the regulation of lysosome trafficking. The mechanisms regulating AP5Z1 degradation are not fully elucidated and other spatacsin interactors identified in our screen may contribute to finely regulate this pathway.

In conclusion, we identify spatacsin as a protein promoting contacts between the ER and lysosomes that regulates the trafficking of lysosomes. We demonstrate that spatacsin, by promoting the lysosomal degradation of AP5Z1, regulates the lysosomal association of AP5Z1 and spastizin. The latter are two motor adaptor proteins with antagonistic role regarding lysosome dynamics. The control of AP5Z1 degradation by spatacsin thus appears as a mechanism to regulate the directionality of lysosome movement, which likely has an impact on lysosome distribution in highly polarized cells such as neurons.

## Experimental procedures

### Mouse models

The Spg11 knockout (*Spg11*^-/-^) model has been previously described (Branchu et al., 2017). This model was generated by inserting two stop codons in exon 32, leading to the loss of expression of spatacsin, and can thus be considered as a functional Spg11 knockout. To obtain this mouse model, we created an intermediate model in which floxed exons 32 to 34 bearing the stop codons were inserted in the reverse orientation in intron 34 (Branchu et al., 2017) (Supplementary Figure 3A). RT-PCR of the transcripts followed by sequencing of brain and spleen samples showed this intermediate model to express the floxed allele, with the splicing of exons 32 to 34 and conservation of the reading frame between exon 31 and exon 35 (Supplementary Figures 3B-C). It was thus equivalent to a functional deletion of exons 32-34 and was named *Spg11^Δ32-34/Δ 32-34^*.

### Antibodies

The antibodies used for immunofluorescence and immunoblotting were rat anti-Lamp1 (clone 1D4B, Development Studies Hybridoma Bank, University of Iowa, USA, deposited by JT August), rabbit anti-V5 (Cat#8137, Sigma), mouse anti-V5 (Cat#Ab27671, Abcam), rat anti-HA (clone 3F10, Cat#11867423001, Merck), rabbit anti-HA (Cat#ab9110, Abcam), rabbit anti-GFP (Cat#6556, Abcam), mouse anti-Myc (clone 9E10, Development Studies Hybridoma Bank), rabbit anti-Stim1 (Cat#5668, Cell Signaling Technology), rabbit anti-cathepsin D (Cat#Ab75852, Abcam), rabbit anti-spatacsin (Cat#16555-1-AP, ProteinTech), rabbit anti-spastizin (Cat#5023, ProSci), rabbit anti-AP5Z1 (Cat#HPA035693, Sigma), mouse anti-clathrin heavy chain (clone 23, Cat#610500, BD Biosciences), rabbit anti-UBR4 (Cat#Ab86738, Abcam), mouse anti p150^Glued^ (Cat#610474, BD Biosciences), mouse anti-α-tubulin (Clone DM1A, Cat#ab7291, Abcam), and mouse anti-actin (Clone C4, Cat#ab3280, Abcam).

The secondary antibodies used for immunofluorescence were purchased from Thermofisher: donkey anti-mouse IgG Alexa 488 (Cat#A21202), goat anti-rabbit IgG Alexa 555 (Cat#A21429) and goat anti-rat IgG Alexa 647 (Cat#A21247). For STED microcopy, the secondary antibodies were goat anti-rabbit IgG STAR 580 (Cat#ST580-1002, Abberior, Göttingen, Germany) and goat anti-mouse IgG STAR 635 (Cat#ST635-1001, Abberior). The secondary antibodies coupled to horseradish peroxidase used for immunoblotting were purchased from Jackson ImmunoResearch (Ely, UK): donkey anti-mouse IgG (Cat#JIR715-035-151) and donkey anti-rabbit IgG (Cat#711-035-152).

### Plasmids

The spastizin-GFP vectors has been previously described (Hanein et al., 2008). Spastizin-HA was obtained by replacing GFP with an HA tag in the spastizin-GFP vector. A codon-optimized vector expressing human spatacsin was generated (Baseclear, Leiden, Netherlands) in a gateway compatible system (Thermofisher). The cDNA was transferred by LR clonase into the pDest-47 vector (Thermofisher), leading to a vector expressing spatacsin-GFP. Deletion of nucleotides 6013 to 6477 of optimized spatacsin cDNA resulted in spatacsin^Δ32-34^. C-terminal fragments of spatacsin (aa 1943-2433) and spatacsin^Δ32-34^ (aa 1943-2226) were amplified by PCR and inserted in the pDest-53 vector (Thermofisher), leading to vectors that expressed GFP-spatacsin-Cter and GFP-spatacsin-Cter^Δ32-34^. Vector expressing AP5Z1-His was obtained from M Slabicki (Słabicki et al., 2010). To generate GFP-AP5Z1 or GFP-mCherry-AP5Z1 constructs, a stop codon was inserted after the final codon of AP5Z1 in the AP5Z1-His vector, and AP5Z1 cDNA was transferred by LR clonase into the pDest-53 or the pDEST-CMV-N-Tandem-mCherry-EGFP vector (Addgene #123216), respectively. The other plasmids used in the study were obtained from other laboratories or Addgene. GFP-Sec61β was obtained from G. Voeltz (Voeltz et al., 2006), AP5Z1-His from M Slabicki (Słabicki et al., 2010, p. 48), reticulon2-V5 from E. Reid (Montenegro et al., 2012)KIF13A-YFP and KIF13A-ST-YFP from C. Delevoye (Delevoye et al., 2014), p150Glued-CC1-mCherry was from T. Schroer (Quintyne et al., 1999), Lamp1-GFP (#16290), Lamp1-mCherry (#45147) and mCherry TRIM46 (#176401) were from Addgene.

### siRNA

The siRNAs used to downregulate Spg11 were either On-target plus siRNAs (Dharmacon), with the sequences CAGCAGAGAGUUACGCCAA (#J-047107-09-0002) and CAGUAUGUGCCGGGAGAUA (#J-047107-12-0002), or from Thermofisher, with the sequence GGUUCUACCAGGCUUCUAUtt (#s103130). The siRNAs used to downregulate AP5Z1 were Silencer Select siRNAs form Thermofisher: GGAGCAGAGUAACCGGAGAtt (#s106997) and UCUGCUCCCGGGUCACUAAtt (#s106999). The siRNAs used to test the role of spatacsin interactors identified by the two-hybrid screen were Silencer Select siRNAs from ThermoFisher and are listed in Supplementary Table 3.

### Isolation of the ER and lysosome-enriched fractions

Isolation of the ER and lysosome-enriched fractions was performed according to previously described protocols with several modifications (Bozidis et al., 2007; Graham, 2000). After killing mice using CO_2_, the brains were immediately extracted and washed with PBS at 4°C. Brains were mechanically dissociated in 250 mM sucrose, 1 mM EDTA, 10 mM Hepes pH 7.4, 1 mM DTT and 25 mM KCl supplemented with a protease inhibitor cocktail (Thermofisher), using a PFTE (polytetrafluoroethylene) pestle attached to a stirrer (Heidolph, Germany) rotating at 500 rpm. Lysates were centrifuged at 800 x g for 5 min and the pellets discarded. The supernatant was centrifuged at 20,000 x g for 10 min and the resulting pellet A retained for lysosome isolation and the supernatant A for ER isolation. For lysosome isolation, pellet A was resuspended in 2 mL of the initial buffer and deposited on 10 ml of 27% Percoll solution in a 15-mL tube. After 90 min of centrifugation at 20,000 x g, the lysosomal fraction was visible close to the bottom of the tube and collected by pipetting. It was then resuspended in the initial buffer and centrifuged at 20,000 x g for 10 min. The pellet was resuspended in sample buffer and analyzed by western blotting.

To isolate the ER, supernatant A was deposited on a gradient of several sucrose solutions prepared in 10 mM Tris pH 7.4 and 0.1 mM EDTA. The sucrose concentrations of the three solutions were from the bottom up: 2 M, 1.5 M, and 1.3 M. The preparation was centrifuged for 70 min at 152,000 x g. After centrifugation, the ER-enriched fraction was found at the phase-limit between the 1.3 M sucrose solution and the 1.5 M sucrose solution. The fraction was collected and resuspended in the initial buffer and centrifuged for 45 min at 152,000 x g. The pellet was resuspended in sample buffer and analyzed by western blotting.

### Mouse embryonic fibroblast cultures

Mouse embryonic fibroblasts were prepared using 14.5 day-old embryos obtained from the breeding of heterozygous (*Spg11*^+/-^ or *Spg11^+/Δ32-34^*) mice as previously described (Boutry et al., 2019). Comparisons between mutant and wild-type fibroblasts were always performed using fibroblasts originating from embryos of the same breeding. All experiments were performed with fibroblasts between passages 4 and 6.

### Transfection of Fibroblasts

Fibroblasts were transfected using the NEON transfection system (Thermofisher) with one pulse of 30 ms at 1350 V, according to manufacturer instructions. Cells (5 x 10^5^) were transfected with 5 μg plasmid and analyzed 24 h later. When we co-transfected a vector expressing a fluorescent protein together with a vector expressing a non-fluorescent protein for live imaging, we imaged cells expressing the fluorescent protein and then fixed the cells afterwards to verify that > 95% of cells expressing the fluorescent marker were also positive for the nonfluorescent protein by immunostaining. For transfection with siRNA, 50 x 10^3^ cells were transfected with 1 pmol siRNA and analyzed after 48 h in culture.

### Primary cultures of neurons and transfections

Mouse cortical neurons were prepared using cortices of 14.5-day-old embryos obtained from the breeding of heterozygous (*Spg11*^+/-^) mice. After a chemical dissociation for 15 min at 37°C (Trypsin 0.05% in Hibernate medium, Thermofisher) and mechanical dissociation, cortices were passed through a 70 microns filter and plated at a density of 100 000 neurons per cm^2^ on pre-coated with Poly-L-Lysine (100 mg/L) 8-well coverslip IBIDI chambers. Neurons were grown in Neurobasal medium supplemented with B27 (ThermoFisher). Neurons were transfected on day 4 of culture. For each well of the IBIDI chamber, we prepared a mix containing 0.6 μl of Lipofectamine 2000 in 15 μl Opti-Mem medium (Thermofisher), with 500 ng of DNA or 1 pmol of siRNA that was added to cultured neurons for 3 hours.

### Chemicals

Lysotracker Blue, Green and Red (Thermofisher) were used at 50 nM for 30 min to stain acidic lysosomes in fibroblasts. DQ-Red-BSA and DQ-Green-BSA (Thermofisher) were added to the culture medium at 2 μg/ml 1 h before imaging and then washed once with culture medium. Texas-Red conjugated dextran (10,000 MW −Thermofisher) was added to the culture medium at 100 μg/ml and the cells incubated for 4 h to allow its internalization by endocytosis and chased for 24 h to stain the lysosomal compartment. The PI3 kinase inhibitor wortmannin (Sigma) was used at 100 nM for 1 h. The proteasome inhibitor MG132 was purchased from Tocris and was added to cells for 16 hours at 15 μM. Bafilomycin (Tocris) was added to cells for 16 hours at 100 nM.

### Immunofluorescence

Cells were fixed in 4% PFA in PBS for 20 min and then permeabilized for 5 min in PBS containing 0.2% v/v Triton X-100. Cells were then blocked for 45 min in PBS with 5% w/v BSA (PBS-BSA) and incubated with primary antibodies in PBS-BSA overnight at 4°C. Cells were washed three times with PBS and incubated with secondary antibodies coupled to fluorophores. After three washes with PBS, glass coverslips were then mounted on glass slides using Prolong Gold antifade reagent (Thermofisher).

### Confocal microscopy

Images of immunofluorescence were acquired using an inverted laser scanning Leica SP8 confocal microscope (Mannheim, Germany) with a 63X objective N.A. 1.40. STED microscopy was performed using a Stedycon device (Abberior) mounted on an inverted Zeiss Imager M2 microscope, with a 100X objective N.A. 1.46.

For live imaging, cells were imaged at 37°C and 5% CO2 using a Leica DMi8 inverted spinning disk confocal microscope equipped with 63X objective N.A. 1.40 and a Hamamatsu Orca flash 4.0 camera. Timelapses of MEFs were acquired to analyze the trajectories of the lysosomes with one image taken every 1 s for 1 min. For timelapses capturing the trajectories of lysosomes in neurons, one image was taken every 500 ms for 30 seconds. Axons were identified by the accumulation of the axon initial segment protein TRIM46 upon transfection of the vector expressing mCherry-TRIM46.

### Electron microscopy

Cultured MEFs were fixed with 2.5% glutaraldehyde in PBS for 2 h at 22 °C. The cells were then post-fixed in 1% osmium tetroxide for 20 min, rinsed in distilled H2O, dehydrated in 50% and 70% ethanol, incubated in 1% uranyl acetate for 30 min, processed in graded dilutions of ethanol (95–100%, 5 min each), and embedded in Epon. Ultrathin (70 nm) sections were cut, stained with uranyl acetate and lead citrate, and analyzed with a JEOL 1200EX II electron microscope at 80 kV.

### Two-Hybrid screen

The yeast two-hybrid screen was performed by Hybrigenics (Paris, France) using an adult human cDNA brain library. The bait was either the complete 1943-2443 domain of human spatacsin or the same domain lacking the amino acids encoded by exons 32 to 34.

### Image analysis

#### Tubular lysosome detection

After detecting lysosomal particles using ICY Spot Detector, the *regionprops* function of the MATLAB Image Processing Toolbox was used to determine the shape characteristics of the particles on the binary images. Tubular lysosomes were defined as follows: circularity < 0.5, eccentricity > 0.9, and a width/length ratio > 4. The selected particles were saved in a new image. To screen the effect of siRNAs on tubular lysosomes, we defined a tubulation index, for which we normalized the number of tubular lysosomes/μm^2^ to a value ranging from 0 to 1, corresponding to the average number of tubular lysosomes/μm^2^ quantified in *Spg11*^-/-^ and *Spg11*^+/+^ fibroblasts analyzed in the same experiment.

#### Lysosome trajectory analysis

Analysis of the movement and trajectories of lysosomes over a minute was performed on timelapse images acquired every second. First, to analyze the trajectory characteristics of round and tubular lysosomes particles, the particles were labeled by hand using the multipoint tool of Fiji software to extract their coordinates. The length of the trajectory and the mean speed were then computed using MATLAB. We then performed automated analysis solely for tubular lysosomes using MATLAB software. The *regionprops* function was used to detect the position of the centroids of tubular lysosomes during the timelapse. Once the coordinates were obtained, they were analyzed using John C.Crocker *track.pro* ‘freeware’ MATLAB function to determine the characteristics of the particle trajectories, considering that the maximum theoretical displacement of a particle between two frames was the approximate size of one tubular lysosome, hence 2.4 μm – 20 pixels. Then, the total distance that each particle traveled was calculated.

#### Measurement of the area of the ER-lysosome overlap

The binarization of lysosomal staining was performed using Spot Detector. The binarization of ER staining was performed using the ImageJ thresholding tool. Once the two binary images were obtained, they were compared using MATLAB and the area of overlap between the two stainings per particle was measured using the *regionprops* function. We calculated a threshold of overlap consistent with a contact site between ER and a lysosome. Considering a 500 nm wide lysosome within 10 nm of distance of a 60 nm wide ER tubule and the fact that the pixel size for the spinning disk at 63X objective is 120 nm, the overlap of the apparent lysosome (740 nm diameter) and the apparent ER tubule is equal to 230 nm, which represents about 30% of the diameter of the lysosomal staining. Therefore, we chose 30% area overlap between the lysosomal and ER stainings as a threshold for the existence of a contact.

#### Analysis of movement directionality using Kymographs

Kymographs of the trajectories of particles along the axons of mouse neurons were obtained in Fiji, using the *Kymograph Builder* plugin. Once the Kymographs were obtained, the proportion of anterograde/ retrograde and stationary movement of lysosomes was identified by hand.

### Image classification by a neural network

Lysosomes of *Spg11*^+/+^ and *Spg11*^-/-^ fibroblasts, as well as *Spg11*^+/+^ fibroblasts transfected with siRNAs, were stained using Texas-Red conjugated dextran. Images were acquired using a spinning disk confocal microscopy, generating an image library.

To train the Spg11 classification model, Tensorflow (https://www.tensorflow.org/?hl=fr) and Scikit-learn (https://scikit-learn.org/stable/) Python libraries were used. Images of lysosomes of *Spg11*^-/-^ (n = 742 cells) and *Spg11^+/+^* (n = 735 cells) MEFs were used as a database. Training and test sets were generated randomly with a test set size of 15% (111 *Spg11*^+/+^ fibroblast images and 112 *Spg11*^-/-^ fibroblast images). Initial images of 921×1024 pixels were resized to 224×224 pixels to reduce input size while retaining consistent information. Data augmentation was performed using flip Tensorflow functions to improve training and artificially increase the number of images. Finally, the image pixel values were normalized between 0 and 1. The transfer learning approach was used to avoid model training from scratch. The VGG16 (Simonyan and Zisserman, 2015) neural network structure was used and downloaded using the Tensorflow_hub library (https://www.tensorflow.org/hub?hl=fr). VGG16 is a convolutional neural network model trained on ImageNet, which is a dataset of over 14 million images belonging to 1,000 classes. The top three layers were excluded and replaced with three other layers: one dense layer of 512 neurons, a dropout layer, and a 64-neuron layer. The final layer was the soft-max layer. The neural network was trained using a NVIDIA GeForce GTX 1050 Ti for 150 epochs, with a starting learning rate at 0.0001 and a batch size at 32. Model evaluation resulted in 79.5% total accuracy on the test set.

The trained model was used to predict the probability of the cell to be considered as a *Spg11*^-/-^ fibroblast for each image of fibroblast transfected with siRNA. For each siRNA, the arithmetic mean of the probability was calculated.

### Protein extraction from cells

MEFs were washed twice with PBS and lysed in 100 mM NaCl, 20 mM Tris pH7.4, 2 mM MgCl2, 1% SDS, and 0.1% Benzonase (Sigma). Samples were centrifuged at 17,000 x g for 15 min and the supernatants recovered as solubilized proteins. The protein concentration was determined using the BCA assay kit (Thermofisher).

### Western Blotting

Protein lysates supplemented with sample buffer (final concentration 80 mM TrisHCl pH 6.8, 10 mM DTT, 2% SDS, 10% glycerol) were separated on 3-8% Tris-acetate or 4-12 Bis-Tris gels (Thermofisher). Proteins were then transferred to PVDF membranes (Merck).

Membranes were then incubated in Ponceau red for 5 min and blocked in PBS-0.05% Tween (PBST) with 5% milk for 45 min. The membranes were incubated with primary antibodies in PBST-5% milk overnight at 4°C. Secondary antibodies were conjugated with HRP (Jackson Lab) and the signals visualized using chemiluminescent substrates (SuperSignal West Dura/Femto; Thermofisher). The chemiluminescent signal was then acquired on Amersham Hyperfilm ECL. Images shown in figures are representative of at least three independent experiments.

### Co-immunoprecipitation

Cells were lysed on ice in 100 mM NaCl, 20 mM Tris pH7.4, 1 mM MgCl_2_, and 0.1% NP40 supplemented with a protease inhibitor cocktail. Samples were centrifuged at 17,000 x g for 15 min at 4°C. Ten percent of the supernatant was retained, supplemented with sample buffer, and was used to monitor protein quantity for the inputs. The remaining 90% of the supernatants was incubated with 10 μL of GFP-trap beads (Chromotek, Germany) for 90 min using a rotating wheel at 4°C. For coimmunoprecipitation of spastizin by KIF13A constructs, beads were washed 4 times in lysis buffer and supplemented with sample buffer with DDT. For co-immunoprecipitation of p150glued by AP5Z1 constructs, beads were washed 4 times in 300 mM NaCl, 20 mM Tris pH7.4, 1 mM MgCl_2_, and 0.1% NP40. Beads and inputs were then analyzed by western blotting.

### Proximity Ligation Assay

MEFs were fixed in PBS containing 4% PFA for 15 min. The Duolink Proximity Ligation Assay (PLA, Sigma) was then performed according to the manufacturer’s instructions. After performing the PLA reaction, we immuno-stained the cells with fluorescent secondary antibodies to detect transfected cells and used the Prolong Gold antifade mounting medium (Thermofisher) instead of the Duolink in situ mounting medium provided with the kit.

### Statistics

Data were analyzed using GraphPad Prism version 9 software. Data normality was assessed using the D’Agostino-Pearson test for sample sets with N > 10. Comparisons of the medians were performed by Mann-Whitney or Kruskall-Wallis tests for non-normally distributed data. Comparisons of the means were performed by unpaired t-tests or ANOVA followed by Sidak’s multiple comparisons test for normally distributed data.

## Acknowledgments

We thank the Phenoparc, Celis, iGenSeq and ICM.quant core facilities of the Paris Brain Institute and Noemi Asfogo for their contributions. This work was supported by “Investissements d’Avenir” program [ANR-10-IAIHU-06] and [ANR-11-INBS-0011] grants and received funding from the European Research Council (European Research Council Starting [grant No 311149] to F.D.). M.B. received a fellowship from the French Ministry of Research (doctoral school ED3C). A.P. received an ARDOC fellowship from the Région Ile de France (grant 17012953; doctoral school ED3C) and a fellowship from the Fondation pour la Recherche Médicale (grant FDT202001010829).

## Supplementary figure legends

**Supplementary Figure 1. Spatacsin is an ER-resident protein**

Live imaging of the ER marker Sec61ß-GFP and lysosome marker Lamp1-mCherry in *Spg11*^+/+^ and *Spg11*^-/-^ MEFs. Note that the absence of spatacsin (*Spg11*^-/-^) did not alter ER morphology. Scale bar: 5 μm.

**Supplementary Figure 2. Spatacsin regulates the dynamics of acidic tubular lysosomes**

A. Live imaging of lysosomes stained with various markers in *Spg11*^+/+^ MEFs. Note that tubular lysosomes were positive for Lamp1, 10 kDa Dextran-Texas Red (TR), Lysotracker, as well as DQ-BSA, indicating that they are acidic and catalytically active compartments. Scale bar: 5 μm.

B. Quantification of the number of tubular lysosomes in *Spg11*^+/+^ and *Spg11*^-/-^ MEFs using the fluorescent markers DQ-BSA-green, Lysotracker-Green, or Dextran-Texas Red. Median and 95% CI, N > 40 cells from three independent experiments. *P < 0.05, Kruskall-Wallis test followed by Dunn’s multiple comparisons test.

C. Quantification of the proportion of lysosomes with an average speed > 0.3 μm/sec according to their shape in wild-type MEFs. Median and 95% CI. **P < 0.01, Mann-Whitney test.

D. Quantification of the average speed of round lysosomes in *Spg11*^+/+^ and *Spg11*^-/-^ MEFs. Median and 95% CI, N = 110 from five independent MEFs. Mann-Whitney test.

**Supplementary Figure 3. Domain of spatacsin encoded by exons 32 to 34 of Spg11 is important for the function of spatacsin**

A. Diagram showing the genomic structure of the mouse *Spg11* gene (top), the targeting vector (middle), and the targeted locus upon excision of the neomycin resistance cassette and action of the Cre-recombinase (bottom). The mutations introduced were c.6052C > T (p.Arg2018*) and c.6061C > T (p.Gln2021*). Scheme adapted from (Branchu et al., 2017).

B. Sequencing of RT-PCR product obtained from the brains of homozygous mice that incorporated the targeting vector, showing the splicing of exons 32, 33, and 34.

C. Scheme representing the mRNA produced in a wild-type mouse, a mouse that incorporated the targeting vector, or after the action of the Cre recombinase. Note that the intermediate model expressing the floxed allele showed splicing of exons 32 to 34 with conservation of the reading frame between exons 31 and 35. It was thus equivalent to a functional deletion of exons 32 to 34, leading to expression of a protein called Spatacsin^Δ32-34^.

D. Western blot showing expression of truncated spatacsin in Spg11^Δ32-34/Δ32-34^ mouse brain. Equal loading was validated by clathrin heavy chain (HC) immunoblotting.

E. Immunostaining of cells expressing the ER marker GFP-Sec61β and V5-tagged spatacsin or V5-tagged spatacsin^Δ32-34^. Cells were immunostained with anti-V5 antibody and the lysosome marker cathepsin D. Scale bar: 5 μm.

F. Western blots of wild-type MEFs transfected with control siRNA or independent siRNA that downregulate spatacsin purchased from Dharmacon (A) or ThermoFisher (B). Lysate of *Spg11*^-/-^ MEFs was used as a negative control. Equal loading was validated by clathrin heavy chain (A) or α-tubulin (B) immunoblotting.

**Supplementary Figure 4. Lysosomal localization of GFP-mCherry-AP5Z1**

Live images of wild-type MEFs expressing GFP-mCherry-AP5Z1 treated with bafilomycin 100 nm for 16h. Note that GFP and mCheryy signals perfectly colocalize. Scale bar: 10 μm

**Supplementary Figure 5. KIF13A and p150^Glued^ contribute to lysosome trafficking**

A. Proportion of tubular lysosomes moving > 1.2 μm over 1 min in wild-type MEFs treated with wortmannin. Median and 95% CI, N = 17 cells from three different independent experiments. **P = 0.014, Mann-Whitney test.

B. Proportion of tubular lysosomes moving > 1.2 μm over 1 min in wild-type MEFs transfected with wild-type KIF13A or mutant KIF13A-ST. Mean and 95% CI, N = 22 cells from three different independent experiments. *P < 0.05, one-way ANOVA, Dunnett’s multiple comparisons test.

C. Proportion of tubular lysosomes moving > 1.2 μm over 1 min in wild-type MEFs transfected with p150^Glued^-CC1-mCherry. Median and 95% CI, N = 65 cells from three different independent experiments. ****P < 0.05, Mann-Whitney test.

**Supplementary Figure 6. AP5Z1 contributes to lysosome directionality**

A. Live image of primary cortical neuron transfected with Lamp1-GFP and mCherry-TRIM46 that labels the axon initial segment, allowing us to identify axons (white arrows). Scale bar: 10 μm.

B. Quantification of the proportion of lysosomes that are moving retrogradely along the axon of *Spg11*^+/+^ and *Spg11*^-/-^ primary mouse neurons. Mean and 95% CI, N = 79 cells from three independent experiments. P=0.1, unpaired t-test.

C. Quantification of the proportion of lysosomes that are moving anterogradely along the axon of *Spg11*^+/+ -^ primary mouse neurons expressing either GFP or GFP-AP5Z1. Mean and 95% CI, N = 32 cells from three independent experiments. P =0.29, unpaired t-test.

D. Quantification of the proportion of lysosomes that are moving retrogradely along the axon of *Spg11*^+/+ -^ primary mouse neurons expressing either GFP or GFP-AP5Z1. Mean and 95% CI, N = 32 cells from three independent experiments. *, P=0.0268, unpaired t-test.

E. Western blot of wild-type MEFs transfected with a control siRNA or siRNA downregulating AP5Z1.

**Supplementary video 1.** Live imaging of wild-type MEF expressing Lamp1-mCherry and Sec61β-GFP. Note the tubular lysosome movement along ER tubular network. One frame was acquired every 750 milliseconds. The video is sped up 4 times. Scale bar: 3 μm.

**Supplementary video 2.** Live imaging of *Spg11*^+/+^ MEF expressing Lamp1-mCherry in green. Detected tubular endolysosomes are highlighted in magenta. One frame was acquired every 750 milliseconds. The video is sped up 4.5 times. Scale bar: 10 μm.

**Supplementary video 3.** Live imaging of *Spgll*^-/-^ MEF expressing Lamp1-mCherry in green. Detected tubular endolysosomes are highlighted in magenta. One frame was acquired every 750 milliseconds. The video is sped up 4.5 times. Scale bar: 10 μm.

