## Supplementary material for "Spatacsin regulates directionality of lysosome trafficking": Supllementary Table1

| Gene name | siRNA#1 | siRNA#2 | Mean |
| --- | --- | --- | --- |
| ENC1 | **0,48** | **0,45** | **0,47** |
| EEf1D | **0,70** | 0,19 | **0,45** |
| EP400 | **0,40** |  | **0,40** |
| ARHGEF6 | **0,50** | **0,28** | **0,39** |
| PSMD2 | **0,37** | **0,39** | **0,38** |
| RNF31 | **0,43** | **0,26** | **0,35** |
| YWHAH | 0,20 | **0,47** | **0,33** |
| ANXA7 | **0,34** | **0,32** | **0,33** |
| ARIH2 | **0,22** | **0,42** | **0,32** |
| HIPK2 | **0,37** | **0,27** | **0,32** |
| EXOC7 | **0,37** | **0,26** | **0,31** |
| COPS4 | **0,42** | 0,19 | **0,31** |
| SMARCE1 | **0,24** | **0,34** | **0,29** |
| VCPIP1 |  | **0,28** | **0,28** |
| LDHA | **0,31** | **0,23** | **0,27** |
| UBR4 | **0,30** | **0,24** | **0,27** |
| MKRN3 | 0,17 | **0,35** | **0,26** |
| KDM5D | **0,26** | **0,25** | **0,25** |
| MAP3K11 | **0,29** | 0,19 | **0,24** |
| SPG7 | **0,23** | **0,23** | **0,23** |
| SPTBN1 | 0,20 | **0,26** | **0,23** |
| ALDOA | 0,20 | **0,26** | **0,23** |
| MYCBP2 | 0,10 | **0,33** | **0,22** |
| USP8 | **0,27** | 0,16 | **0,21** |
| TRIP12 | 0,20 | **0,23** | **0,21** |
| USP14 | **0,29** | 0,13 | 0,21 |
| MOAP1 | 0,16 | **0,24** | 0,20 |
| NEFL | **0,23** | 0,17 | 0,20 |
| PDS5B | 0,20 | 0,17 | 0,18 |
| PIK3CB | 0,10 | **0,26** | 0,18 |
| CARS2 | **0,23** | 0,13 | 0,18 |
| IFT172 | **0,23** | 0,10 | 0,16 |
| VPS8 | 0,13 | 0,19 | 0,16 |
| TIAM1 | 0,17 | 0,13 | 0,15 |
| TP53BP1 | 0,10 | 0,20 | 0,15 |
| PBXIP1 | 0,13 | 0,10 | 0,11 |
| BLZF1 | 0,06 | **0,23** | 0,14 |
| FRY | 0,17 | 0,10 | 0,13 |
| IFI30 | 0,07 | 0,20 | 0,13 |
| DMAP1 | 0,10 | 0,16 | 0,13 |
| SPARCL1 | 0,13 | 0,11 | 0,12 |
| KCTD9 | 0,19 | 0,03 | 0,11 |
| CLU | 0,10 | 0,10 | 0,10 |
| KCNAB2 | 0,10 | 0,07 | 0,08 |
| TTC8 | 0,03 | 0,03 | 0,03 |

**Supplementary table 1 :** Unbiased analysis of the effect of siRNA downregulating genes encoding putative binding partners of domain of spatacsin encoded by exons 32-34 of *SPG11*. The scores represent the probablility of phenocopying *Spg11^-/-^* MEFs (see methods). Bold indicates genes that are at least as efficient as three independent Spg11 siRNA (SPG11#1 : 0.36 ; SPG11#2 : 0.21 ; SPG11#3 : 0.26).
