## Supplementary Table2 for "Spatacsin regulates directionality of lysosome trafficking"

| Gene name | siRNA#1 | siRNA#2 | Mean |
| --- | --- | --- | --- |
| PSMD2 | **0.08** | **0.37** | **0.22** |
| UBR4 | **0.17** | **0.31** | **0.24** |
| ARHGEF6 | **0.42** | **0.14** | **0.28** |
| EEf1D | **0.39** | **0.28** | **0.34** |
| IFT172 | **0.31** | **0.40** | **0.36** |
| FRY | **0.34** | **0.46** | **0.40** |
| BLZF1 | **0.42** | **0.42** | **0.42** |
| USP8 | **0.38** | **0.47** | **0.42** |
| TRIP12 | 0.56 | **0.35** | **0.46** |
| KDM5D | **0.36** | 0.60 | **0.48** |
| ENC1 | 0.71 | **0.38** | **0.55** |
| RNF31 | 0.67 | **0.48** | 0.57 |
| COPS4 | **0.48** | 0.7 | 0.59 |
| VCPIP1 |  | 0.59 | 0.59 |
| EXOC7 | 0.72 | **0.47** | 0.60 |
| KCTD9 | 0.62 | 0.62 | 0.62 |
| TTC8 | 0.72 | **0.52** | 0.62 |
| SMARCE1 | 0.68 | 0.63 | 0.66 |
| SPARCL1 | 0.73 | 0.58 | 0.66 |
| NEFL | 0.76 | 0.59 | 0.68 |
| TIAM1 | 0.87 | 0.58 | 0.72 |
| KCNAB2 | 0.85 | 0.73 | 0.79 |
| USP14 | 0.66 | 0.92 | 0.79 |
| TP53BP1 | 0.88 | 0.73 | 0.81 |
| LDHA | 0.64 | 1 | 0.82 |
| PDS5B | 0.82 | 0.83 | 0.82 |
| PBXIP1 | **0.45** | 1.29 | 0.87 |
| CARS2 | 0.67 | 1.10 | 0.88 |
| HIPK2 | 0.82 | 0.95 | 0.88 |
| YWHAH | 0.69 | 1.09 | 0.89 |
| EP400 | 0.89 |  | 0.89 |
| MYCBP2 | 1.07 | 0.80 | 0.93 |
| CLU | 1.22 | 0.66 | 0.94 |
| IFI30 | 1.13 | 0.82 | 0.98 |
| SPTBN1 | 1.09 | 0.97 | 1.03 |
| MKRN3 | 1.09 | 0.98 | 1.04 |
| DMAP1 | 1.41 | 0.68 | 1.05 |
| ANXA7 | 1.01 | 1.10 | 1.06 |
| ARIH2 | 1.12 | 1.10 | 1.11 |
| ALDOA | 1.08 | 1.19 | 1.13 |
| MOAP1 | 1.10 | 1.15 | 1.13 |
| PIK3CB | 1.60 | 0.66 | 1.13 |
| VPS8 | 1.20 | 1.08 | 1.14 |
| MAP3K11 | 1.46 | 0.89 | 1.18 |
| SPG7 | 0.92 | 1.55 | 1.23 |

**Supplementary Table 2** : Analysis of the proportion of tubular lysosomes in control MEFs transfected with siRNA downregulating genes encoding putative binding partners of domain of spatacsin encoded by exons 32-34 of SPG11. The scores represent the normalized number of tubules (score = 1 for control MEFS, score = 0 for *Spg11^-/-^* MEFS). Bold indicates genes that are at least as efficient as three independent Spg11 siRNA (SPG11#1 : 0.55 ; SPG11#2 :0.43 ; SPG11#3 : 0.50) to decrease the proportion of tubular lysosomes.
