## Supplementary Figures for "Spatacsin regulates directionality of lysosome trafficking"

Pierga et al. Supplementary Figure 1

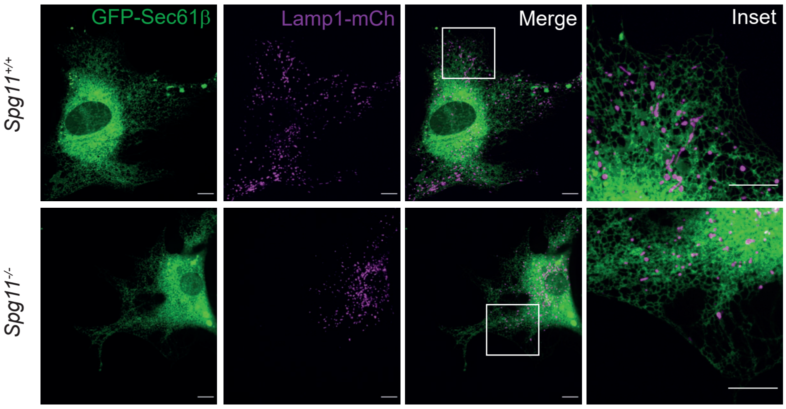

Pierga et al. Supplementary Figure 2

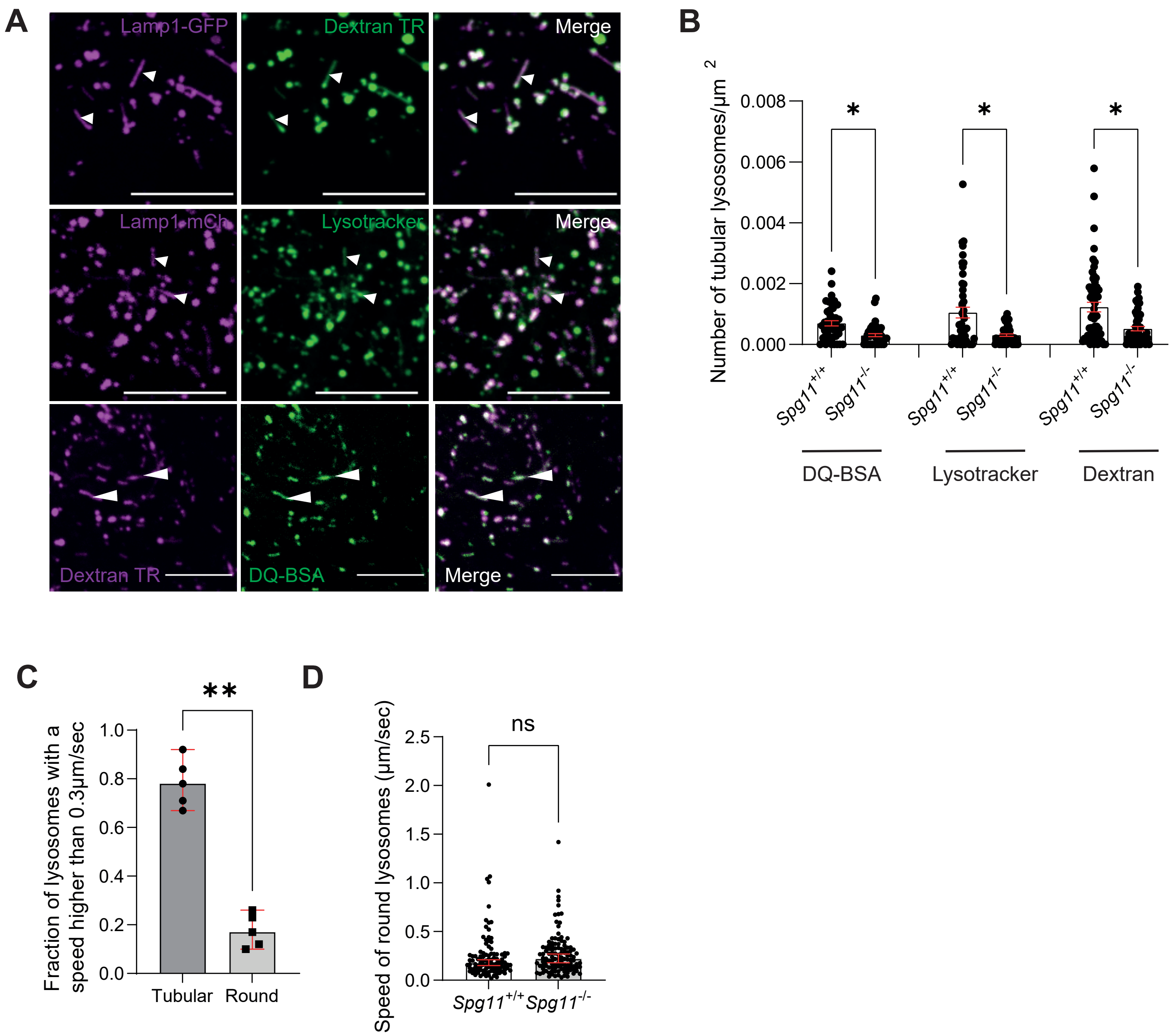

Pierga et al. Supplementary Figure 3

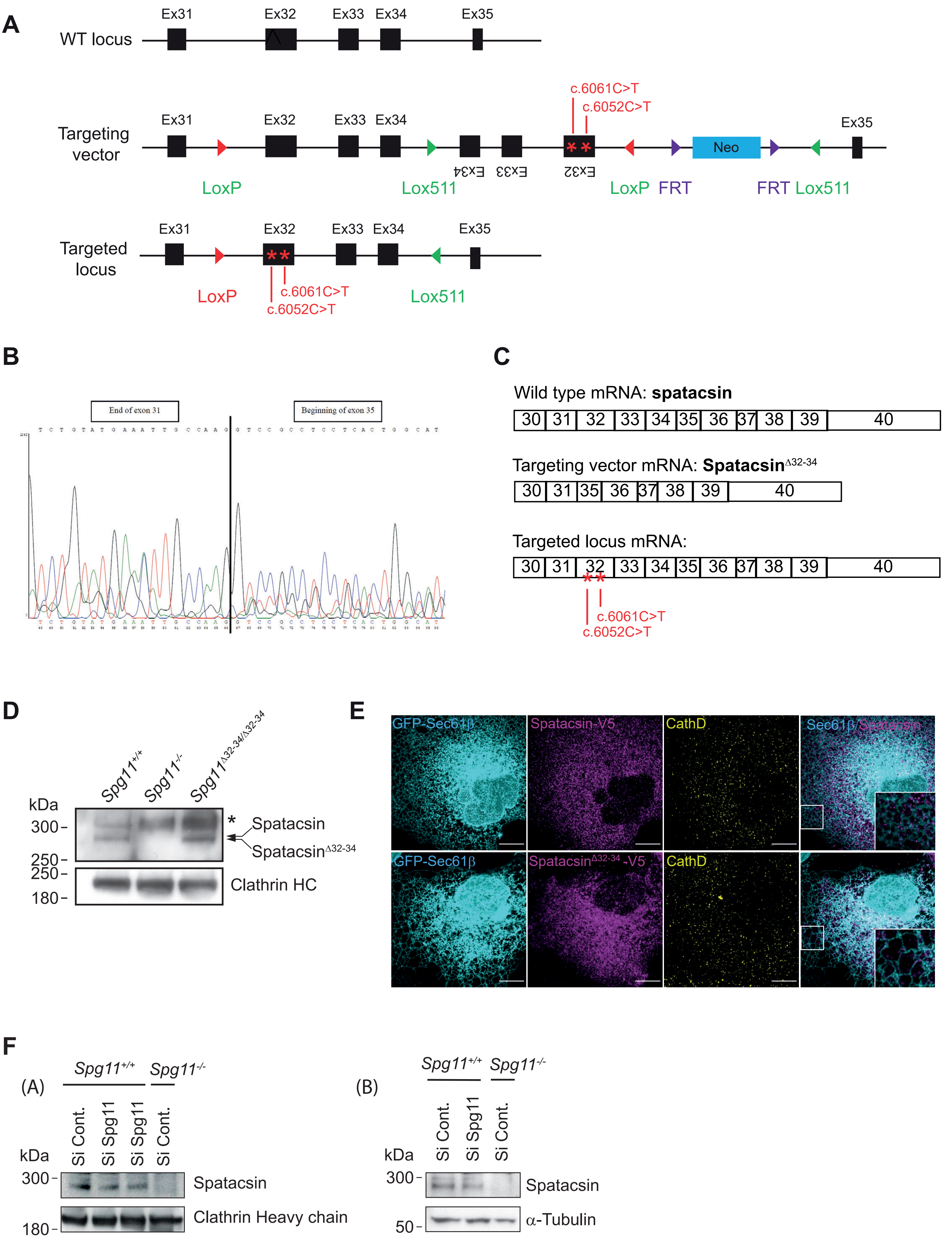

Pierga et al. Supplementary Figure 4

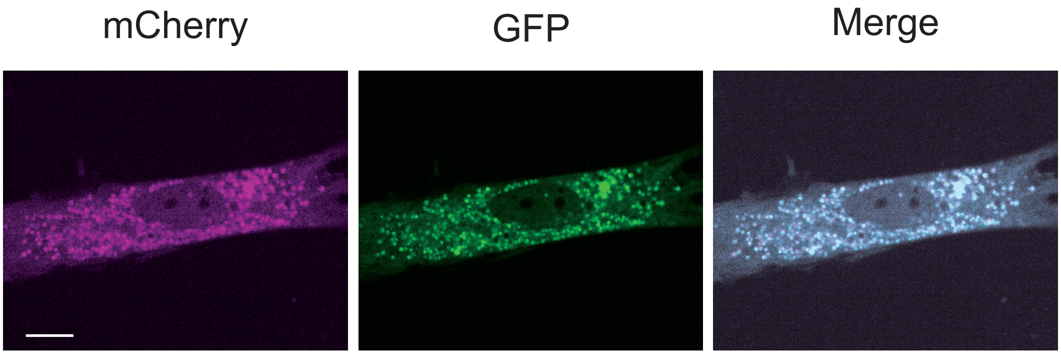

Pierga et al. Supplementary Figure 5

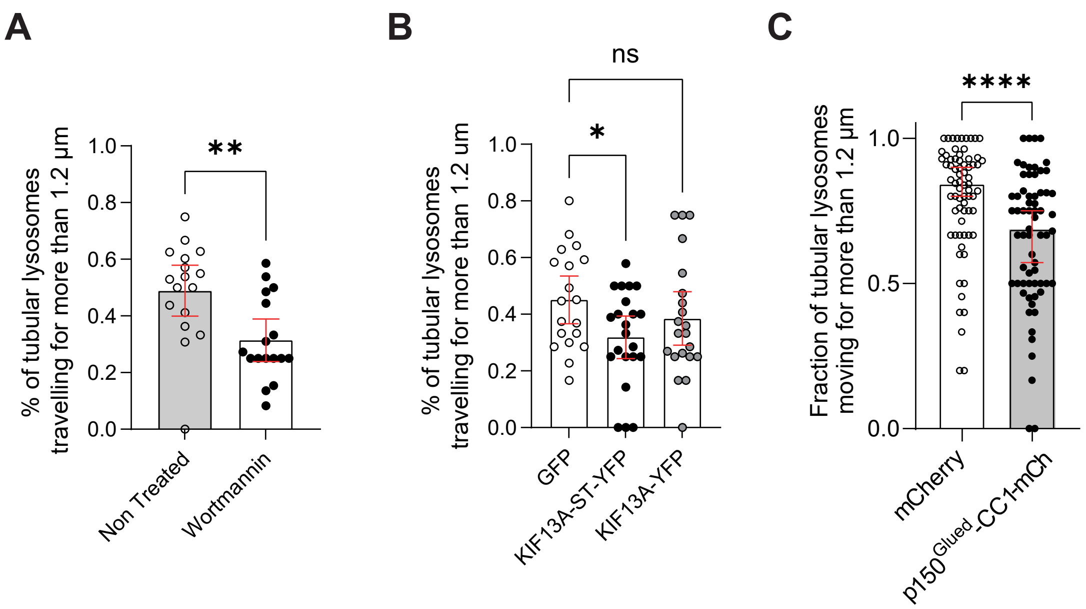

Pierga et al. Supplementary Figure 6

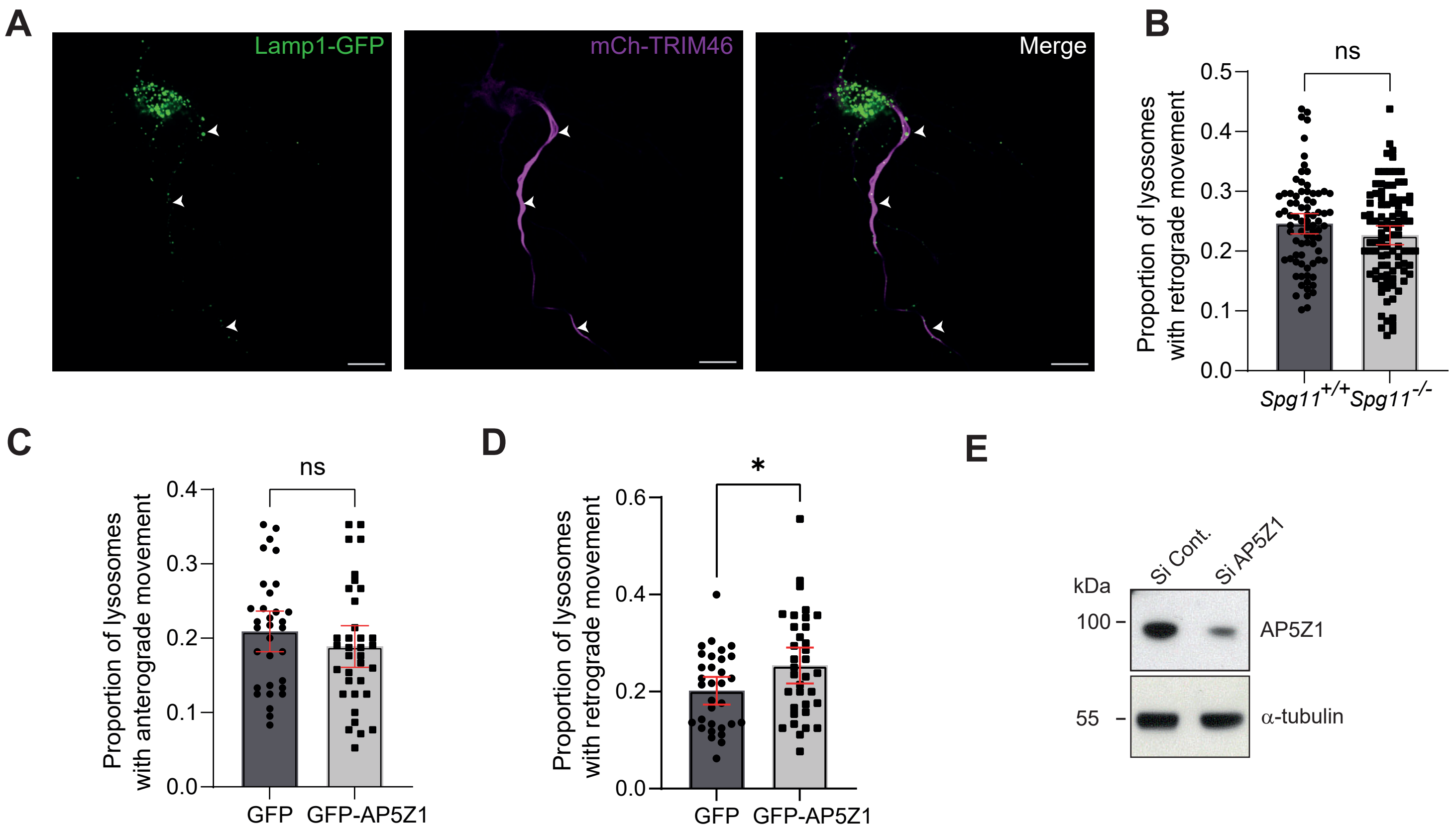
